# Control of poly(A)-tail length and translation in vertebrate oocytes and early embryos

**DOI:** 10.1101/2023.10.18.562922

**Authors:** Kehui Xiang, Jimmy Ly, David P. Bartel

## Abstract

During oocyte maturation and early embryogenesis, changes in mRNA poly(A)-tail lengths strongly influence translation, but how these tail-length changes are orchestrated has been unclear. Here, we performed tail-length and translational profiling of mRNA reporter libraries (each with millions of 3ʹ-UTR sequence variants) in frog oocytes and embryos, and fish embryos. We found that the UUUUA element, together with the polyadenylation signal (AAUAAA or AUUAAA), specifies cytoplasmic polyadenylation, and identified contextual features that modulate the activity of both elements. In maturing oocytes, this tail lengthening occurs against a backdrop of global deadenylation and the action of C-rich elements that specify tail-length-independent translational repression. In embryos, cytoplasmic polyadenylation becomes more permissive, and additional elements specify waves of stage-specific deadenylation. Together, these findings largely explain the complex tapestry of tail-length changes observed in early frog and fish development, with strong evidence of conservation in both mice and humans.

**Highlights:**

1. UUUUA, modulated by contextual features, specifies most cytoplasmic polyadenylation
2. Stage-specific sequence motifs drive waves of tail-length shortening in embryos
3. UUUUA and C-rich motifs can direct tail-length-independent translational repression
4. Tail-length control is conserved in oocytes of frogs, mice, and humans

## INTRODUCTION

In vertebrate animals, transcription ceases when oocytes become fully grown and stays quiescent throughout oocyte maturation, fertilization, and early embryogenesis, until activation of the zygotic genome.^1–3^ To progress through these transcriptionally silent developmental stages, oocytes and early embryos rely on post-transcriptional control of a stockpile of mRNAs that had been maternally deposited. In zebrafish (*Danio rerio*) and frogs (*Xenopus laevis*), most of the maternally deposited mRNAs are stable until zygotic transcription ensues during the mid-blastula transition (MBT).^4,5^ Precise temporal regulation of translation from these mRNAs is crucial for the oocyte-to-embryo transition and early embryogenesis.^6–8^

Much of this regulation occurs through changes in mRNA poly(A)-tail lengths. In oocytes and early embryos of flies, fish, frogs, and mice, mRNA translational efficiency and poly(A)-tail length are strongly coupled, such that mRNAs with long poly(A) tails are translated much more efficiently than those with short tails.^9–12^ In frog prophase I-arrested oocytes, most mRNAs have short poly(A) tails and are translationally repressed.^13^ As these oocytes mature and complete meiosis I, a group of mRNAs, including *mos* and *ccnb1*, which encode proteins essential for meiosis re-entry, undergo cytoplasmic polyadenylation.^14,15^ This lengthening of their poly(A) tails causes translational activation of these mRNAs, thereby promoting oocyte maturation,^16–18^ a process conserved in mammals.^19–21^

Cytoplasmic polyadenylation requires at least two sequence elements within mRNA 3ʹ UTRs: a polyadenylation signal (PAS) and a cytoplasmic polyadenylation element (CPE).^22,23^ The PAS has the consensus sequence AAUAAA (with the top alternative variant being AUUAAA), the same motif used for nuclear 3ʹ-end cleavage and polyadenylation. It is recognized by the cleavage and polyadenylation specificity factor (CPSF), a protein complex shared with nuclear polyadenylation.^24^ The CPE is recognized by the CPE-binding protein (CPEB1).^25^ The most common CPE motifs are reported to be UUUUAU and UUUUAAU, although additional, nonconsensus motifs (UUUUACU, UUUUCAU, UUUUCCA, UUUUAAAU, UUUUAAGU) have also been implicated.^21,23^ However, these sequence motifs are each identified from only a few endogenous mRNAs, relying on Northern blots for poly(A) tail-length measurements, without the benefit of transcriptome-wide analyses. Recent studies have used high-throughput sequencing methods to measure mRNA poly(A)-tail lengths transcriptome-wide during oocyte maturation in frogs, mice, and humans.^26–29^ However, little progress has been made to define sequence motifs that modulate tail lengths or to quantitively determine the effects of such motifs, in part due to 1) the limited sequence space represented by a few thousand endogenous mRNAs, and 2) difficulty in designating causal sequence motifs from many co-occurring sequences within 3ʹ UTRs.

Cytoplasmic polyadenylation is also essential for early embryogenesis,^30,31^ but the primary sequence elements that drive tail lengthening in the embryo are less well understood. In frog embryos, a few mRNAs are reported to employ motifs that differ from the CPE used in oocytes, including eCPE^32^ and C-rich motifs,^33^ but the prevalence of these motifs in tail-lengthening of the transcriptome is unknown. Besides polyadenylation, mRNAs of early embryos are also subject to deadenylation through elements recognized by RNA-binding proteins or microRNAs.^34–38^ Indeed, in fly embryos, poly(A)-tail lengthening appears to occur in a sequence-independent manner, with specificity primarily imposed by tail shortening.^10,39^ In flies, fish, and frogs, shortening of poly(A) tails is essential for translational repression and then clearance of most maternal mRNAs upon onset of zygotic transcription.^36–38^ How embryos control poly(A)-tail lengths of different mRNAs through the two arms of tail length-modulating processes in successive developmental stages is not well understood, with much to be learned about the 3ʹ-UTR regulatory code for poly(A)-tail length over the course of oocyte maturation and early embryonic development.

Here, we generated diverse mRNA libraries containing millions of 3ʹ-UTR sequence variants and examined the tail lengths of library molecules after injection into developing oocytes or embryos. These analyses defined a core CPE for cytoplasmic polyadenylation and revealed contextual features that modulate its effect on mRNA poly(A)-tail lengths. Our analyses also identified elements that direct deadenylation at specific developmental stages and revealed co-existing tail-length dependent and independent translational regulatory regimes.

## RESULTS

### A core CPE motif controls cytoplasmic polyadenylation

To acquire information useful for identifying the 3ʹ-UTR features that specify cytoplasmic polyadenylation during oocyte maturation, we generated a large library of mRNA variants (Figure 1A), injected this mRNA library into prophase I-arrested frog oocytes, and both sequenced the 3ʹ UTR and measured the poly(A)-tail length for individual variants over the course of oocyte maturation (Figure 1B). Each member of the mRNA library contained the Nanoluc coding region, followed by a 3ʹ UTR containing a 60-nucleotide (nt) random-sequence region, designed to match the mean nucleotide sequence composition of frog 3ʹ UTRs (30% A, 19% C, 19% G, and 32% U). This random-sequence region was followed by a segment from the last 21 nt of *X. laevis mos.L* 3ʹ UTR, which contained the PAS and was followed by a 35-nt poly(A) tail. We refer to this mRNA library as N60-PAS*^mos^*.

**Figure 1.**
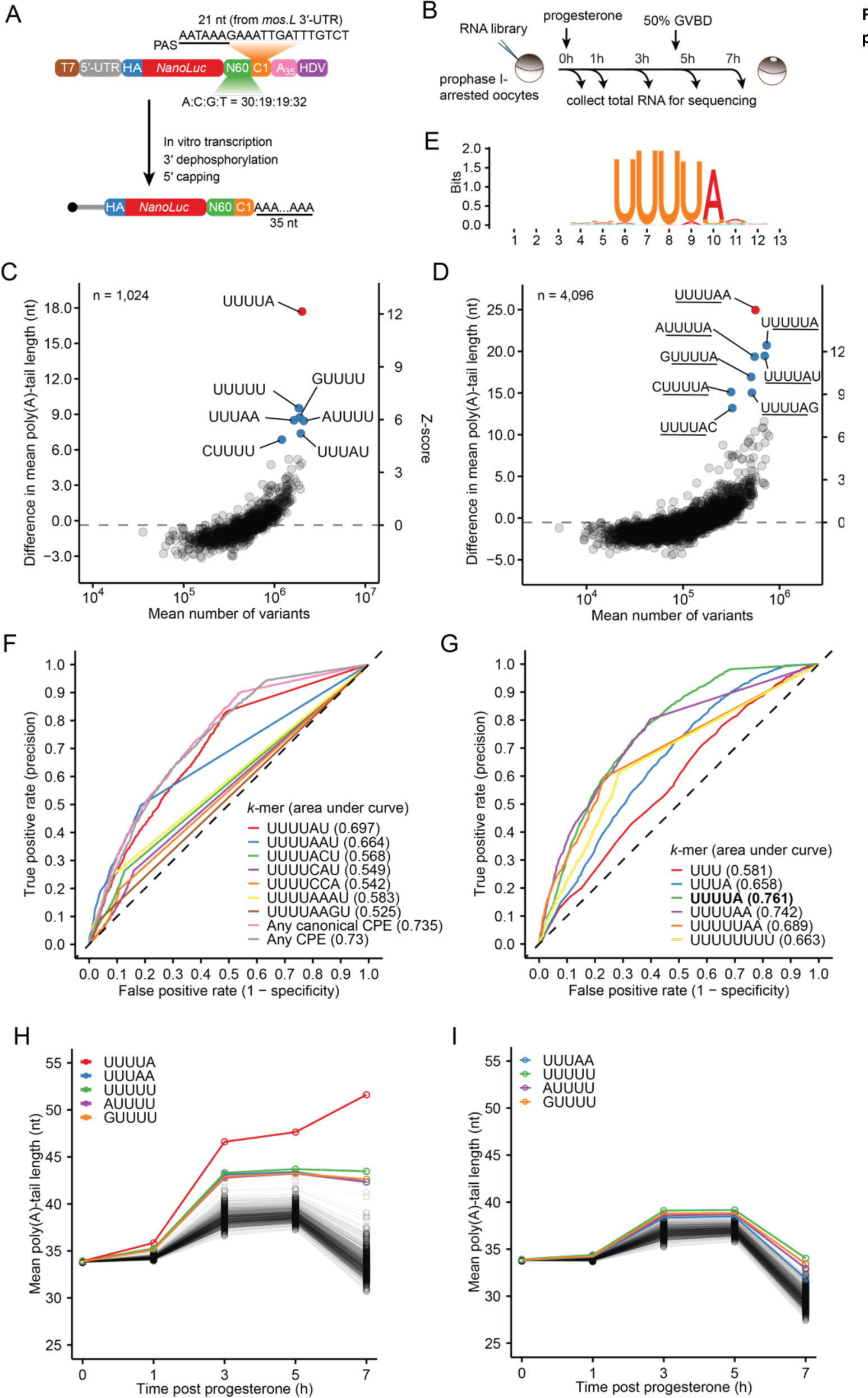
A core CPE controls cytoplasmic polyadenylation in frog oocytes. (A) Schematic of the N60-PAS*^mos^* mRNA library used for injection. (B) Experimental scheme for mRNA library injection and sample collection during frog oocyte maturation. (C) Tail-length changes associated with each 5-mer in the 3ʹ UTRs of the N60-PAS*^mos^* library, comparing between 0 and 7 h post progesterone treatment. Plotted for each 5-mer are differences in mean poly(A)-tail lengths observed for mRNAs with that 5-mer. Circles represent 5-mers associated with significant differences in mean poly(A)-tail lengths (Welch’s *t*-test against the global average, adjusted *P* < 0.01), whereas squares represent 5-mers associated with non-significant changes. (D) As in (C), but for 6-mers. (E) Sequence logo generated from the 180 8-mers most strongly associated with lengthening of mRNA tails in the N60-PAS*^mos^* library, comparing between 0 and 7 h post progesterone treatment. (F) ROC curves testing the ability of previously defined CPE motifs to classify endogenous mRNAs as subject to cytoplasmic polyadenylation during frog oocyte maturation. A curve is shown for each of the indicated CPE motifs, including the two canonical CPE motifs (UUUUAU and UUUUAAU), as well as for the indicated combinations of motifs. (G) ROC curves testing the ability of the best 3-, 4-, 5-, 6-, 7-, and 8-mer to classify endogenous mRNAs as subject to cytoplasmic polyadenylation during frog oocyte maturation. (H) The effects of 5-mers on tail lengths of mRNAs in the N60-PAS*^mos^* library during frog oocyte maturation. Plotted for each 5-mer are the mean tail lengths of mRNAs containing that 5-mer. (I) As in panel H, except all mRNA variants that contained the UUUUA motif were excluded.

Analysis of these library molecules was expected to have several advantages over analysis of endogenous mRNAs. First, for each library molecule, the variable sequence was presented within a uniform context, which avoided potentially confounding effects of either different mRNA lengths, different initial poly(A)-tail lengths, different 5ʹ-UTR or coding region sequences, or different nuclear or cytoplasmic histories. Second, a larger number of different mRNAs could be examined, with this number limited only by the depth of sequencing (tens of millions) rather than by the number of different expressed mRNAs (tens of thousands). Third, and most importantly, the use of reporters that presented each *k*-mer (sequence motif of length *k*) within a diversity of non-biological sequences enabled causal relationships between *k-*mers and tail length to be confidently established. In contrast, biological mRNAs contain *k-*mers that tend to co-occur with each other, which can confound establishment of causality. For example, if a motif that specifies mRNA localization and a motif that specifies polyadenylation tend to co-occur with each other within natural 3ʹ UTRs, both might correlate with the extension of natural mRNA tails, even though only one of them functions to specify polyadenylation.

As oocytes underwent maturation, the median tail length of the mRNA library gradually lengthened until after 5 h, the time of germinal vesical breakdown (GVBD) followed by meiosis I, after which point the median tail length substantially shortened (Figure S1A). By 7 h post progesterone treatment, 30% of mRNAs had poly(A) tails shorter than 20 nt (Figure S1A), supporting the idea that short-tailed mRNAs are stable in maturing frog oocytes.^40,41^

To identify sequence motifs that mediated cytoplasmic polyadenylation, we evaluated all 5-nt *k*-mers, determining for each 5-mer the average poly(A)-tail length 7 h after progesterone treatment and comparing this length to that observed for motif-containing mRNAs in the uninjected sample. The greatest difference in tail length was observed for the UUUUA 5-mer; variants with this motif had average tail lengths that were 17.7 nt longer in the matured oocytes compared to the uninjected sample (Figure 1C). In contrast, the next six strongest 5-mers were associated with weaker tail lengthening (6.9–9.5 nt). Moreover, they each resembled a partial UUUUA motif that if extended by a U on the 5ʹ terminus or an A on the 3ʹ terminus would form the full UUUUA motif.

Analyses of shorter and longer *k*-mers confirmed the strong association of UUUUA with tail lengthening. When all 4-mers were examined, UUUU and UUUA—the two 4-mers that constitute the UUUUA motif, clearly separated from the others (Figure S1B). Similarly, among all 6-mers, the eight that were extensions of the UUUUA motif were associated with the longest tail-length differences (Figure 1D).

These results indicated that UUUUA is the CPE of frog oocytes. This conclusion was somewhat in contrast to previous results, in that previously identified CPEs (UUUUAU, UUUUAAU, UUUUACU, UUUUCAU, UUUUCCA, UUUUAAAU, UUUUAAGU) are all longer than 5 nt, such that those that contain UUUUA are each extended by one or more additional nucleotides at the 3ʹ end. To eliminate the spillover signal from a stronger motif to a weaker motif, we re-examined the effects of 6-mers by iteratively removing all mRNA variants containing the 6-mer with the largest association with tail-lengthening and then re-calculating the average tail lengths of all remaining 6-mers. After excluding the top two 6-mers (UUUUAA and UUUUAU), other 6-mer motifs, including UUUUUA, AUUUUA, UUUUAG, and UUUUAC, which would not have previously been recognized as a CPE, were associated with substantial tail lengthening (Figure S1C), thereby supporting the designation of the shorter, UUUUA motif as the CPE. Moreover, a search for sequence logos from even longer, 8-mers that were associated with elongated poly(A) tails in matured oocytes identified only a single consensus logo, which closely resembled the UUUUA motif, with relatively little information contributed by flanking nucleotides (Figure 1E). Therefore, we refer to this core UUUUA motif as the CPE of frog oocytes. The close match between the UUUUA CPE and the binding site for human CPEB1 in vitro^42^ and for fly CPEB1 in vivo^43^ suggested that these results extend to diverse species, and further suggested that binding of CPEB1 protein confers the specificity of CPE recognition without help from additional factors.

To examine if the CPE identified in our mRNA library might also act in the context of endogenous mRNAs, we measured tail lengths of mRNAs collected alongside the mRNA library during frog oocyte maturation. Globally, tail-length distribution of endogenous mRNAs followed a similar trend to that of the injected mRNA library, with its median value increasing before GVBD and decreasing afterward (Figure S1D). Because *Xenopus laevis* 3ʹ UTRs were poorly annotated, we used our sequencing data to identify mRNA poly(A) sites and thereby infer 3ʹ-UTR sequences of mRNAs expressed in these oocytes. With these improved annotations and the ability to measure median tail lengths for individual mRNA sequences (which was possible for endogenous mRNAs, due to their relatively lower sequence diversity, but not for mRNAs of our library), we evaluated the performance of different *k-*mers in classifying endogenous mRNA substrates for cytoplasmic polyadenylation using receiver operating characteristic (ROC) curves. In this analysis, UUUUA outperformed each of the previously reported CPEs (Figure 1F–G). Moreover, it had a greater area under the curve (AUC value) than any combination of previously reported CPEs (Figure 1F–G). Indeed, among all *k-*mers with lengths of 3–8 nt, the UUUUA CPE had the greatest AUC value (Figure 1G), regardless of the tail-length difference threshold used for categorizing the true positives (Figure S1E).

Because cytoplasmic polyadenylation is thought to occur in early and late phases during frog oocyte maturation,^44^ we returned to our analysis of library molecules to examine the dynamics of tail-length changes. When mean poly(A)-tail lengths of mRNAs containing each of the 5-mers were plotted during the course of maturation, the CPE was the motif consistently associated with the longest tails (Figure 1H). From 1–3 h CPE-dependent tail lengthening was accompanied by a modest overall increase in tail lengths, whereas after 5 h (upon GVBD), CPE-dependent tail lengthening opposed an overall decrease in tail lengths. This latter behavior was consistent with previous reports that endogenous mRNAs lacking a CPE undergo non-sequence-specific deadenylation in mature oocytes.^45,46^

During the 1–3 h interval, some U-rich motifs were also associated with modest tail-lengthening that did not fully abate when all CPE-containing variants were excluded from the analysis, which indicated that some preferential tail lengthening could be achieved without a perfect match to the CPE (Figure 1I). These U-rich motifs each matched four contiguous nucleotides of the UUUUA CPE, which suggested that their association might have been due to the promiscuous binding of CPEB1 to CPE-like motifs. Nonetheless, these degenerate CPEs were typically not sufficient to overcome the global deadenylation occurring during the 5–7 h time interval, such that after GVBD, the UUUUA CPE was the only motif to robustly support net polyadenylation among 5-mers (Figure 1I) or even 8-mers (Figure S1F).

Despite distinct global tail-length trajectories before and after GVBD, motif-associated tail-length changes in the early (1–3 h) and the late (5–7 h) stages of polyadenylation were highly correlative (Figure S1G), suggesting that sequence-specific regulation of poly(A)-tail length is consistent throughout frog oocyte maturation. Moreover, no 5-mers had evidence of maturation-stage-specific synergistic effects with the CPE in polyadenylation (Figure S1H). Notably, we did not observe substantial tail lengthening in our library for several motifs that were previously implicated in cytoplasmic polyadenylation, including Musashi,^47^ TCS,^48^ and Pumilio^44^ motifs (Figure S1F). Because our global analyses did not detect an effect of these motifs on tail lengths, the previous observations might be attributed to effects on CPE context for the previously analyzed mRNAs.^49^

Taken together, our results indicate that sequence-specific poly(A) tail-length control during frog oocyte maturation is primarily mediated by cytoplasmic polyadenylation that requires a CPE: UUUUA. This specific tail lengthening occurs against a backdrop of non-sequence-specific deadenylation after GVBD.

### CPE context explains differences in cytoplasmic polyadenylation

Although the average tail length of CPE-containing mRNAs in our injected library increased by 17.7 nt in oocytes 7 h after progesterone treatment, the tail lengths for individual variants varied substantially (Figure S2A). We sought to identify contextual features that contributed to this variation, in a search enabled by the large diversity of 3ʹ-UTR sequences in our mRNA library.

When examining the CPE position, we found that the magnitude of cytoplasmic polyadenylation increased as the CPE approached the PAS (Figure 2A). This relationship was almost linear until the distance between the CPE and the PAS reached approximately 6 nt, at which point it dipped slightly, before dramatically declining at 2 nt. We attribute the dip at 6 nt to inhibitory pairing between the CPE and the PAS, and the dramatic decline at 2 nt to a steric clash between CPEB1 and CPSF when the CPE and the PAS motifs were too close. Similar positional relationships with polyadenylation were observed for partial or extended CPE motifs of different lengths (Figure 2A) and for many CPE-like motifs (Figure S2B). The apparent effect of CPE–PAS proximity was somewhat at odds with a report that varying the distance between the CPE and the PAS within a 26 nt range does not affect cytoplasmic polyadenylation,^50^ a difference attributable to the lower resolution of the previous study, which examined a limited number of mRNAs resolved on northern blots.

**Figure 2.**
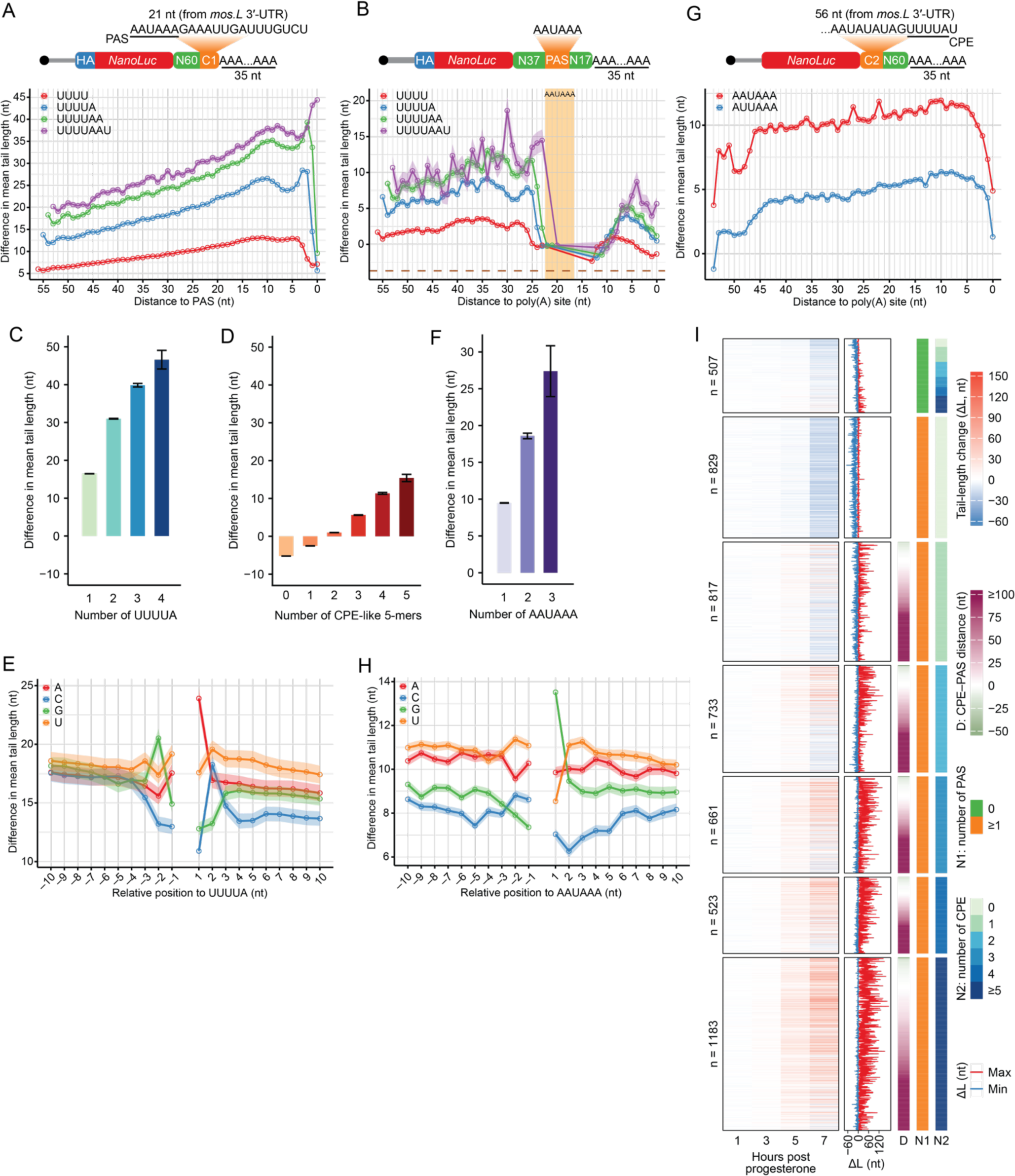
Context of the CPE and the PAS influences cytoplasmic polyadenylation in frog oocytes (A) The effect of the distance between the CPE and the PAS for cytoplasmic polyadenylation. Plotted for each of the four motifs is the difference in mean poly(A)-tail length observed for mRNA variants of the N60-PAS*^mos^* library (schematic at the top) containing that motif at each position along the variable region of the 3ʹ UTR (plotted as distance to the PAS), comparing between 0 and 7 h after progesterone treatment. Shaded areas along the lines indicate standard error of the difference between means. (B) The effect of the relative position between the CPE and the PAS for cytoplasmic polyadenylation. Plotted for each of the four motifs is the difference in mean poly(A)-tail length observed for mRNA variants of the N37-PAS-N17 library containing that motif at each position along the 3ʹ UTR (plotted as distance to the 3ʹ end). The shaded region indicates the location of the PAS. The dashed brown line indicates the difference in mean poly(A)-tail length of all mRNA variants in the library. Otherwise, this panel is as in (A). (C) The influence of the number of the CPEs on cytoplasmic polyadenylation. Plotted is the difference in mean poly(A)-tail length of mRNA variants in the N60-PAS*^mos^* library containing the indicated numbers of CPEs, comparing between 0 and 7 h after progesterone treatment. Error bars indicate standard error of the difference between means. (D) The influence of the number of CPE-like motifs on cytoplasmic polyadenylation. This panel is as D, but for CPE-like motifs, UUUUU, UGUUU, GUUUU, UUUGU, and AUUUU, which were the top five motifs associated with increased poly(A)-tail length after excluding all variants containing UUUUA. (E) The effect of CPE flanking nucleotides on cytoplasmic polyadenylation. Plotted is the difference in mean poly(A)-tail length observed for mRNA variants in the N60-PAS*^mos^* library containing one CPE and the indicated nucleotide at the indicated position relative to the CPE, comparing between 0 and 7 h after progesterone treatment. Shaded areas along the lines indicate standard error of the difference between means. (F) The influence of the number of PASs on cytoplasmic polyadenylation. Plotted is the difference in mean poly(A)-tail length of mRNA variants in the CPE*^mos^*-N60 library containing the indicated numbers of PASs, comparing between 0 and 5 h after progesterone treatment. Error bars indicate standard error of the difference between means. (G) The effect of the distance between the PAS and the mRNA 3ʹ end for cytoplasmic polyadenylation. Plotted is the difference in mean poly(A)-tail length of mRNA variants in the CPE*^mos^*-N60 library containing the indicated PAS motif at each position along the variable region of the 3ʹ UTR (plotted as distance to the 3ʹ end); otherwise as in (A). (H) The effect of PAS flanking nucleotides on cytoplasmic polyadenylation. Plotted is the difference in mean poly(A)-tail length observed for mRNA variants in the CPE*^mos^*-N60 library containing one PAS (AAUAAA) and the indicated nucleotide at the indicated position relative to the PAS, comparing between 0 and 5 h after progesterone treatment. Shaded areas along the lines indicate standard error of the difference between means. (I) The association of 3ʹ-UTR sequence features with poly(A) tail-length changes of endogenous mRNAs of frog oocytes. The heatmaps on the left compare median poly(A)-tail lengths of frog oocyte mRNAs collected at the indicated time after progesterone treatment to those in oocytes not treated with progesterone, with each row representing a unique 3ʹ UTR of an mRNA with a defined poly(A) site. Also indicted for each of these UTRs is the CPE–PAS distance and the minimal and maximal tail-length change over the 7 h course of treatment. UTRs are grouped based on the presence of a canonical PAS (within 150 nt of the 3ʹ end), and number of CPEs (within 1000 nt of the 3ʹ end). Only UTRs with poly(A) sites that had more than 50 poly(A) tags in all datasets were included in this analysis. For UTRs that contained more than one PAS, the one closest to the 3ʹ end was used. For UTRs that contained more than one CPE, the one closest to the PAS was used to calculate the CPE–PAS distance.

To interrogate the positional effect of the CPE downstream of the PAS, we repeated our experiments with another mRNA library we refer to as N37-PAS-N17, in which the PAS was flanked by random-sequence regions on both sides (Figure 2B). Upstream of the PAS, the CPE had a positional relationship resembling that observed for the N60-PAS*^mos^* library (Figure 2B), which indicated that the interplay between these two motifs is not restricted to mRNAs modeled from frog *mos.L* mRNA. Downstream of the PAS, the extent of polyadenylation gradually increased until the distance between the PAS and the CPE reached 6 nt, after which it started to decline (Figure 2B). These results showed that the relative position of the CPE on either side of the PAS influences cytoplasmic polyadenylation.

Besides the relative position, the number of the CPEs within the 3ʹ UTR also impacted cytoplasmic polyadenylation, with more CPEs leading to larger tail-length increases (Figure 2C). In most scenarios, two CPEs on the same mRNA had less effect on polyadenylation than the summed effect of individual CPEs at the same positions, the exceptions being when one CPE was very close to the PAS (distance < 2 nt) or when both CPEs were far away from the PAS (distance > 30), in which case the two CPEs appeared to have synergistic effects (Figure S2C). For mRNA variants without any CPE, multiple copies of CPE-like motifs were able to support some polyadenylation, but approximately five CPE-like motifs were required to approach the tail-length extension imparted by a single CPE (Figure 2C–D, S2D).

Next, we asked if nucleotides flanking the CPE affected polyadenylation. To this end, we focused our analysis on mRNA variants that contained only one CPE. Overall, a U-rich context promoted stronger polyadenylation (Figure 2E), presumably due to secondary CPE-like motifs, which are generally U-rich (Figure S2B). By contrast, a C-rich context was generally associated with the weaker polyadenylation (Figure 2E). The nucleotide identity at –2, –1, +1, and +2 positions had the strongest average impact on polyadenylation, with polyadenylation enhanced by A and U at both the –1 and +1 positions, G at the –2 position, and C as well as U at the +2 position. The impact of these nucleotides at positions flanking the CPE explained why some of the longer motifs were originally designated as CPEs. Nevertheless, the mean contributions to polyadenylation by these flanking nucleotides were much weaker than those by the first and the last nucleotides of the 5-mer CPE (Figure S2E–F), which supported designation of the 5-mer motif as the core CPE.

The flanking dinucleotide effects on polyadenylation could not be explained by the predicted structural accessibility of the CPE (Figure S2G). Although structural accessibility has been implicated in modulating cytoplasmic polyadenylation for select frog endogenous mRNAs^49^ and predicted accessibility correlates well with efficacy of other regulatory sites, such as those of microRNAs,^51,52^ we did not find meaningful correlations between tail lengths and predicted accessibility of either the CPE or the PAS (Figure S2H). Nonetheless, tail length modestly correlated with the predicted minimum free energy of 3ʹ UTR folding even after GC-content was matched (Figure S2H), implying that folding of the 3ʹ UTR might negatively impact cytoplasmic polyadenylation.

Next, we assessed the function of PAS and its contextual features during cytoplasmic polyadenylation. Because our N60-PAS*^mos^* and N37-PAS-N17 libraries both contained a pre-configured canonical PAS motif, AAUAAA, we repeated our experiments with a third mRNA reporter library designated CPE*^mos^*-N60, in which the 3ʹ UTR of each mRNA had a 56-nt fragment from *X. laevis mos.L* that contained a CPE but lacked a PAS, followed by a 60-nt random-sequence region. After injection into oocytes, the 6-mer motif from the random-sequence region associated with the strongest tail lengthening was the canonical PAS motif, AAUAAA, and the next six 6-mers were AUUAAA and five other near matches to the top motif (Figure S2I). Interestingly, like the CPE motifs, more PAS motifs resulted in greater tail-length increases (Figure 2F). Moreover, for single PAS motifs, the magnitude of cytoplasmic polyadenylation increased as the distance between the PAS and the 3ʹ end became shorter, until ∼5 nt, at which point it began to drop (Figure 2G). This positional effect occurring for distances of 45–5 nt defied a diminishing tail-lengthening effect that would be expected due to the increased distance between the CPE and the PAS (Figure 2A), indicating that the distance between the PAS and the 3ʹ end has an even a stronger influence on cytoplasmic polyadenylation than the distance between the PAS and the CPE.

The flanking nucleotides of the PAS also impacted cytoplasmic polyadenylation. At most flanking positions, a U or A enhanced polyadenylation (Figure 2H). The main exception was at the +1 position, where a G strongly increased polyadenylation, whereas a C or U were more detrimental than at most other positions (Figure 2H).

To examine if principles elucidated from analyses of mRNA reporter libraries also applied to endogenous mRNAs, we evaluated tail-length changes of endogenous mRNAs during frog oocyte maturation as a function of the number of CPEs, the presence of a PAS, and the distance between the CPE and the most proximal PAS (Figure 2I). Consistent with the results from analyses of mRNA reporters, larger tail-length increases correlated with more CPEs, the presence of a PAS, and CPE–PAS proximity. For 3ʹ UTRs with only 1 or 2 CPEs, tail lengthening was minimal if these CPEs were distant from the PAS (≥ 100 nt). Interestingly, mRNAs that had no canonical PAS (neither AAUAAA nor AAUAAA) within the last 150 nt of their 3ʹ UTR but did have ≥5 CPEs frequently underwent detectible tail-length extension, albeit to a much lesser degree than those that had a PAS, which implied that binding of multiple CPEB1s might partially compensate for weak association between CPSF and a non-canonical PAS in promoting cytoplasmic polyadenylation.

In summary, we found that cytoplasmic polyadenylation that occurs during frog oocyte maturation is influenced by various contextual features, including nucleotides flanking the CPE and the PAS, the number of CPEs and PASs, the CPE–PAS distance, the PAS–poly(A)-site distance, and to a lesser extent, the structural accessibility of the 3ʹ UTR, with every indication that these contribute to the control of endogenous mRNA poly(A)-tail lengths.

### The CPE and deadenylation motifs successively control tail length during frog early embryogenesis

To extend our investigation beyond matured oocytes to early embryogenesis, we injected our first two mRNA libraries (N60-PAS*^mos^* and N37-PAS-N17), as well as another library, referred to as N60, in which the 3ʹ UTR was composed of 60 nt of random-sequence RNA (Figure S3C), into in vitro fertilized frog eggs at the one-cell stage (stage 1) and harvested total RNA samples at successive developmental stages until gastrulation (stage 12) (Figure 3A). For each library, we calculated the average tail lengths of mRNAs with each 5-mer at each time point to identify sequence-specific regulation of tail lengths (Figure 3B, S3D–E). When comparing adjacent time points, we also calculated changes in mean tail lengths for mRNAs with each 8-mer (Figure 3B).

**Figure 3.**
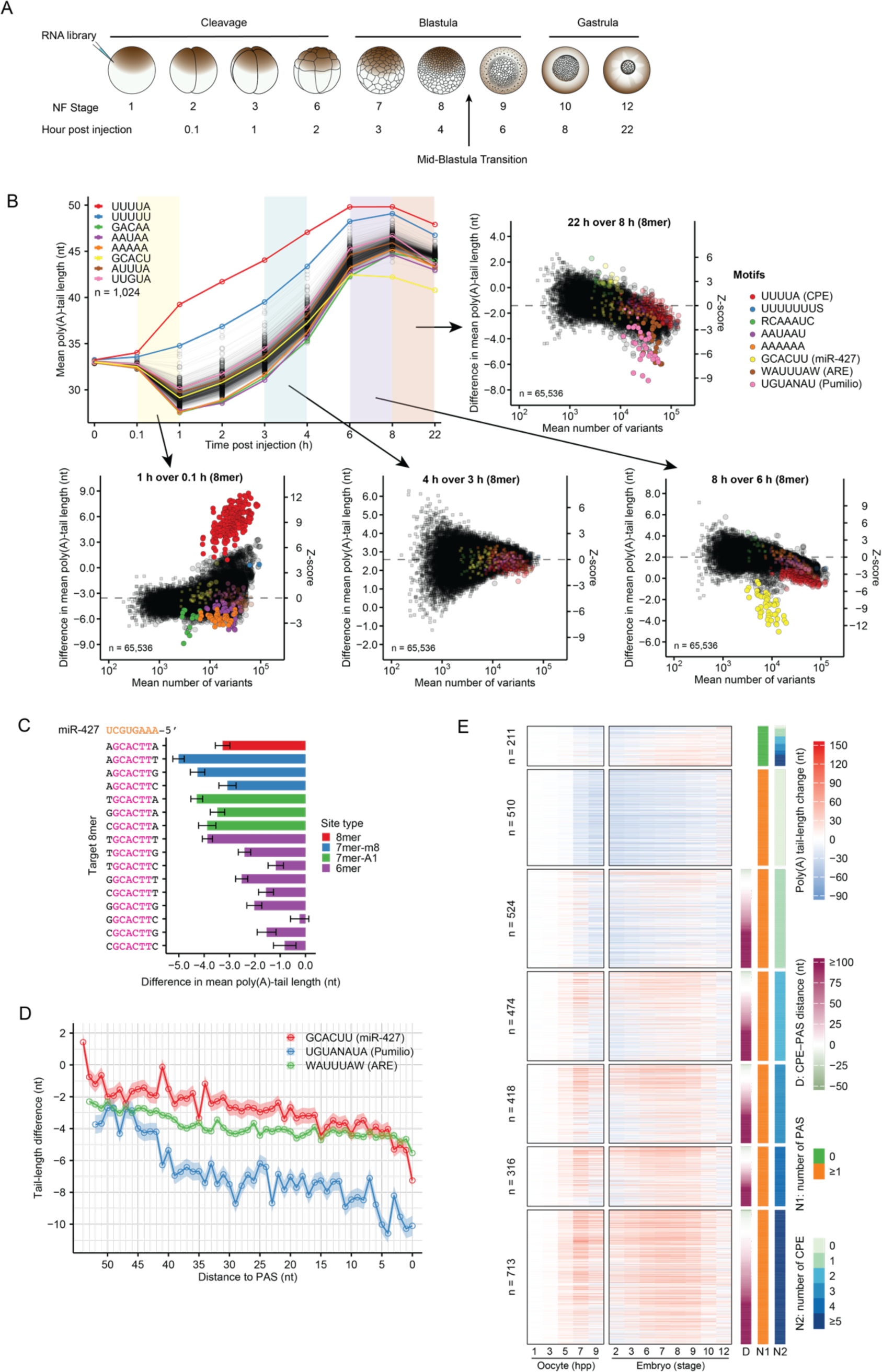
The CPE and deadenylation motifs successively control tail length during frog early embryogenesis (A) Experimental scheme for mRNA library injection and sample collection at different frog embryonic stages. Embryo illustration created based on drawings from digital images on Xenbase.^90,91^ (B) Effects of different 3ʹ-UTR sequence motifs on poly(A)-tail lengths of mRNAs of the N60-PAS*^mos^* library at successive stages of frog early embryogenesis. Plotted on the top left, for each 5-mer over the course of early embryonic development, are mean poly(A)-tail lengths of mRNA variants in the library containing that 5-mer. Shaded areas highlight adjacent stages between which poly(A) tail-length differences are assessed in the peripheral scatter plots. Each scatter plot shows mean tail-length changes associated with each 8-mer in the 3ʹ UTRs of mRNA variants in the library, plotted as in Figure 1C, except circles represent 8-mers with significant differences in mean poly(A)-tail lengths (Welch’s *t*-test against the global average, adjusted *P* < 0.01). Circles representing notable motifs are colored (key, S represents C or G; R represents A or G; W represents A or U; N represents A, C, G, or U). (C) Effects of nucleotides flanking the miRNA seed match on miR-427-directed deadenylation. Shown are the mean tail-length changes of mRNA variants in the N60-PAS*^mos^* library for 16 different 8-mer sites, each centered on the 6-mer miR-427 seed match (brown), comparing between 6 h post injection (stage 9) and 8 h post injection (stage 10). Results for 8-mers that match canonical 7- and 8-nt target sites are colored (key). Error bars indicate standard error of the difference between means. (D) Positional effects of select motifs that promote deadenylation. Plotted for each motif are the differences in mean poly(A)-tail length of mRNA variants in the N60-PAS*^mos^* library at varying distances to the PAS, comparing between 6 h post injection (stage 9) and 8 h post injection (stage 10) for the miR-427 target site (GCACUU) and between 8 h post injection (stage 10) and 22 h post injection (stage 12) for the PBE (UGUANAUA) and ARE (WAUUUAW). Shaded areas along the lines indicate standard error of the difference between means. (E) The effect of 3ʹ-UTR sequence features on poly(A) tail-length changes for frog mRNAs in oocytes and early embryos. The main heatmaps on the left were split between oocytes at different times during maturation (hpp, hours post progesterone) and embryos at different developmental stages; otherwise as in Figure 2I.

In the early period of embryogenesis (before stage 3, within 1 h post injection [hpi]), the median tail lengths of each mRNA library rapidly shortened (Figure S3A–C). Nonetheless, mRNAs with both a CPE and a PAS tended to undergo tail lengthening (Figure 3B, Figure S3D–E). These results resembled those observed during the late stages of oocyte maturation (Figure 1H), suggesting that early embryos inherited the tail-length regulatory regime observed in matured eggs. At this period, deadenylation appeared to be the default, which could be countered by the CPE, the PAS, or long stretches of U (Figure 3B, S3F–H). Although a U_12_ motif has been implicated as an embryonic cytoplasmic polyadenylation element,^32^ we found this motif did not appear to promote polyadenylation as effectively as the CPE (Figure S3I). We also found several sequence motifs that were associated with greater-than-average tail-length shortening (Figure 3B, S3F–H). One of them, with a consensus sequence AAUAAU, was identified in analyses of all three libraries. This motif, although rich in A and U, differed from the canonical AU-rich element (ARE) AUUUA that is known to promote deadenylation in frog embryos at later stages^34^ (Figure 3B), implying that an unknown protein or mechanism might be involved.

During late cleavage and early blastula stages (1–4 hpi; stage 3–8), the median tail length of the N60-PAS*^mos^* library gradually increased while that of the N37-PAS-N17 and the N60 libraries did not (Figure S3A–C). This increase was attributed to promiscuous cytoplasmic polyadenylation mediated by CPE-like motifs within the constant region of the N60-PAS*^mos^* library (Figure S3J), an effect not observed in maturing oocytes, implying that either the extent of global deadenylation decreased at these embryonic stages or cytoplasmic polyadenylation became more permissive. During this period, we did not find any sequence motifs other than the CPE that strongly associated with tail-length changes in any of the three libraries (Figure 3B, S3D–E). The deadenylation occurring before and during this period resulted in a substantial accumulation of tail-less mRNAs in the N37-PAS-N17 and the N60 libraries (Figure S3B–C), supporting the idea that poly(A) tails are not required for mRNA stability during frog early embryonic development.^34^ In contrast, these deadenylated mRNAs and other short-tailed mRNAs appeared to be degraded after 4 hpi (stage 8), causing large increases in the median tail lengths of all three libraries (Figure 3B, S3A–E). These results highlighted a functional transition of poly(A)-tail length—from enhancing translation efficiency to enhancing mRNA stability—during frog embryonic development.

During and immediately after the mid-blastula transition (4–8 hpi; stages 8–10), mRNAs that contained microRNA-427 (miR-427) sites were specifically deadenylated. This concurred with an accumulation of miR-427 after zygotic genome activation,^38^ analogously to the deadenylation directed by miR-430 in zebrafish embryos.^36^ As expected, different miR-427 site types and flanking nucleotides had distinct influences on the extent of deadenylation (Figure 3C), with the effect of flanking sequences corresponding well to relative binding affinities measured for miRNA–Ago2 complexes and their targets.^52^ The miR-427 sites also tended to be more effective when located closer to the mRNA 3ʹ ends (Figure 3D).

In gastrulating embryos, miR-427-associtated deadenylation dampened, giving way to preferential deadenylation of mRNAs that contained either an ARE or a Pumilio-Binding Element (PBE) (Figure 3B), both of which also had 3ʹ-end positional efficacy resembling that of miR-427 target sites (Figure 3D). These motif-specific deadenylation activities coincided with the timing of increased expression of the ARE-binding protein Zfp36 (TTP in mammals) and Pumilio proteins,^53^ the latter of which have been shown essential for embryonic development in flies^54^ and mice,^55^ but their roles during gastrulation of frog embryos had not been reported.

To investigate whether our library-elucidated sequence motifs play similar roles in the temporal regulation of tail lengths of endogenous mRNAs, we measured tail lengths of endogenous mRNAs at the stages of frog embryonic development at which our injected libraries had been sampled. At early stages, when transcription was still silent (stages 2–8), tail-length extension required the CPE and resembled that observed in matured oocytes, although dependence on the PAS was more relaxed, with many mRNAs containing ≥2 CPEs but no canonical PAS undergoing tail lengthening (Figure 3E). Most mRNAs whose tails were extended earlier, during oocyte maturation, either maintained their long tails or underwent further tail-length extension (Figure 3E). Overall, mRNA tail lengths at each stage during this period of embryonic development correlated well with each other (Pearson correlation coefficient, *R*_p_ = 0.82–0.98, Figure S3K) and also well with those in matured oocytes (9 h post progesterone, *R*_p_ = 0.7–0.86, Figure S3K).

The effects of individual motifs on endogenous mRNA tail lengths were difficult to detect due to different starting tail lengths and various confounding motifs within 3ʹ UTRs of different mRNAs. Nevertheless, when these factors were matched between cohorts of mRNAs either containing or not containing a motif, we observed statistically significant support for impact of most elements identified from analyses of our libraries, at embryonic stages matching those at which we observed impact on the reporter libraries (Figure S3L–P). The only exception was the preferential deadenylation of PBE-containing transcripts, observed for reporter mRNAs between stages 10–12 but not for endogenous mRNAs (Figures 3B and S3Q). This lack of signal for PBE activity in endogenous mRNAs might have been due to newly transcribed mRNAs with long tails confounding the analysis. These results illustrate the advantage of using injected mRNAs to identify regulatory elements that are otherwise difficult to identify within endogenous mRNAs.

In summary, we found that frog early embryos achieved temporal regulation of mRNA tail lengths by combining more permissive, oocyte-inherited, CPE-mediated cytoplasmic polyadenylation and stage-specific, motif-driven deadenylation.

### Early fish embryos use a more permissive CPE and undergo more global polyadenylation

To examine tail-length regulation in fish, we injected the N37-PAS-N17 and N60 libraries into zebrafish 1-cell embryos and monitored poly(A)-tail lengths as embryos developed to the stage of gastrulation (shield stage, Figure 4A). The tail-length control in zebrafish embryos resembled that of frog embryos in some respects. For example, most injected mRNAs underwent deadenylation within the first hour, with the strongest effect observed for those that contained poly(A) motifs, AAUAAU, or AREs (Figure 4B–C, S4A–B). Moreover, after zygotic genome activation, miR-430, the fish homolog of frog miR-427, directed deadenylation of its target mRNAs^36^ (Figure 4B–C, S4A–B).

**Figure 4.**
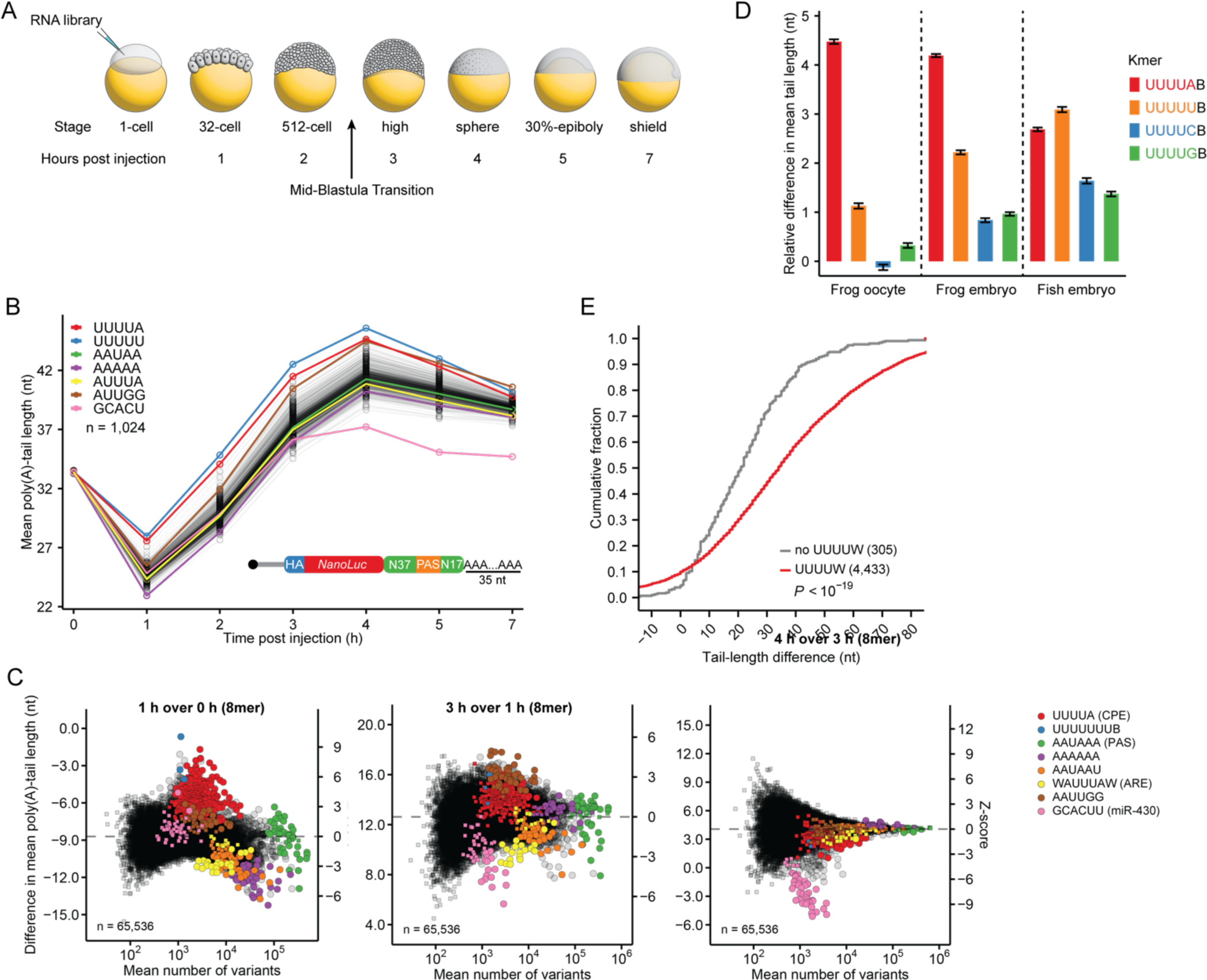
Early fish embryos use a more permissive CPE and undergo more global polyadenylation (A) Experimental scheme for mRNA library injection and sample collection at successive embryonic stages of zebrafish. Illustration of embryos created from drawings in the reference.^80^ (B) The effects of 5-mers on tail lengths of mRNAs in the N37-PAS-N17 library during zebrafish early embryogenesis. Plotted for each 5-mer are the mean tail lengths of mRNAs containing that 5-mer within 3ʹ UTRs. The inset shows the schematic of the N37-PAS-N17 library used for injection. (C) Mean tail-length changes associated with each 8-mer in the 3ʹ UTRs of mRNA variants in the N37-PAS-N17 library, plotted as in Figure 3B. (D) Effects of the CPE and related motifs on poly(A)-tail length in different biological contexts. Shown are mean tail-length changes of mRNA variants in the N37-PAS-N17 library containing the indicated sequence motifs (plotting changes relative to mean tail lengths of all variants; B represents C, G, or U), comparing between 0 and 7 h post progesterone treatment in frog oocytes, between 0.1 and 1 h post injection in frog embryos, and between 0 and 1 h post injection in zebrafish embryos (right). Error bars indicate standard error of the difference between means. (E) Cumulative distributions of tail-length changes for endogenous zebrafish mRNAs with 3ʹ UTRs that contained a fish CPE (UUUUW) and those that did not contain any CPE, comparing between 1-cell stage and high stage embryos. The numbers of mRNAs in each group are listed in parentheses; *P* value, Mann–Whitney U test.

Despite these similarities, poly(A) tail-length control observed in zebrafish embryos differed from that of frog embryos in important respects. First, the 5-mer associated with the longest tails was UUUUU, not UUUUA (Figure 4B). Unlike in frog oocytes and embryos, where cytoplasmic polyadenylation strongly preferred an A at the last position of the CPE, a U was slightly favored for the CPE in zebrafish embryos (Figure 4D). Thus, we designate UUUUW (where W = A or U) as the core CPE in zebrafish embryos.

Second, after the initial deadenylation phase, most injected mRNAs were polyadenylated. This polyadenylation was not driven by any specific short sequence motif, because the mean tail lengths of mRNAs with nearly every 5-mer increased to a similar extent (12 nt on average) during this period (from the 32-cell stage to the high stage). Moreover, the tail-length increase occurred for both the N37-PAS-N17 and N60 libraries (Figure 4B, S4A) and persisted even when all mRNA variants that contained either a UUUU or a canonical PAS (AWUAAA) motif were excluded (Figure S4C), showing that cytoplasmic polyadenylation during these stages did not require either a CPE or a PAS. Nonetheless, mRNAs containing the CPE or an AAUUGG motif underwent slightly more polyadenylation (Figure 4C, S4B).

These observations extended to endogenous mRNAs of fish embryos, in that most of these mRNAs (> 90%) underwent tail lengthening before the high stage, regardless of whether they had a CPE or not, although those that contained a UUUUW CPE had substantially longer tail extensions (Figure 4E). Like for our injected libraries, we also observed stronger tail-lengthening for endogenous mRNAs that contained an AAUUGG element (Figure S4D), the mechanism of which merits further investigation.

### Frog oocytes and embryos modulate translation using both tail-length-dependent and tail-length-independent mechanisms

Because poly(A)-tail length can strongly influence mRNA translational efficiency (TE) in frog oocytes and early embryos,^9,26,41^ we asked how 3ʹ UTR sequences that control poly(A)-tail length impact translation in these developmental contexts. To this end, we injected the N60-PAS*^mos^* library into oocytes and used a sucrose cushion to pellet translating ribosomes from lysates prepared at different developmental stages (Figure 5A). Enrichment for a *k*-mer in the mRNA of the pellet compared to that in the input served as a proxy for its effect on TE.

**Figure 5.**
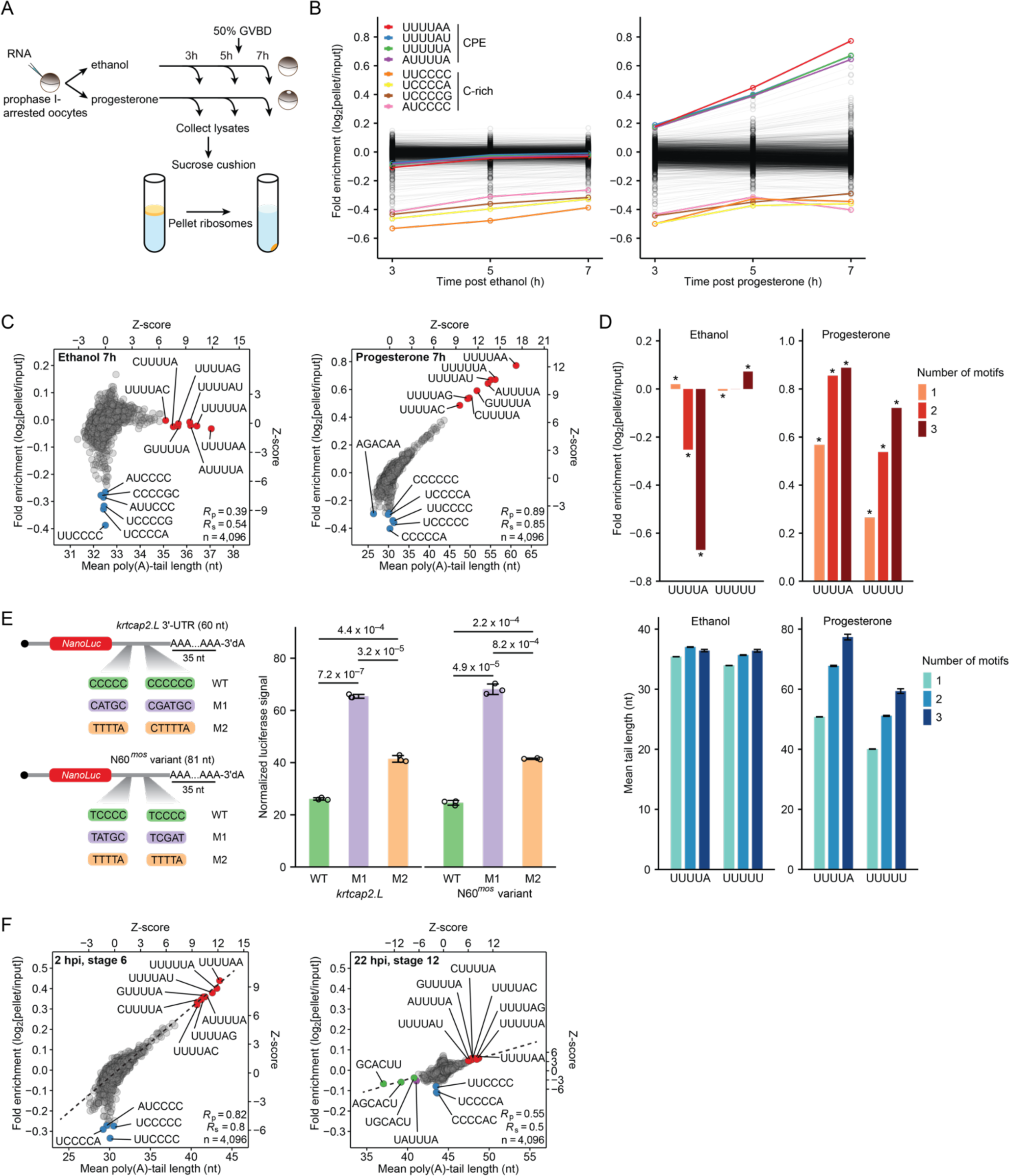
Translation in frog oocytes and early embryos is controlled by both tail-length-dependent and tail-length-independent mechanisms. (A) Experimental scheme for translational profiling of frog oocyte mRNAs by ribosome pelleting with a sucrose cushion. For each mRNA, translational efficiency is measured as the fold enrichment observed between the pellet and the input. (B) The effects of 6-mer motifs on TE of mRNAs in the N60-PAS*^mos^* library during frog oocyte maturation. Results are shown for control prophase I-arrested oocytes that had been treated with ethanol (left) and for maturing oocytes that had been treated with progesterone (right). (C) Translational and tail-length regulation by 3ʹ-UTR sequence motifs in frog oocytes. Shown for each 6-mer in the 3ʹ UTRs of the N60-PAS*^mos^* library is the mean translational efficiency (as measured by fold enrichment in pelleted ribosomes) observed for mRNAs with that 6-mer, plotted as a function of mean poly(A)-tail length of these mRNAs. Results are shown for prophase I-arrested oocytes (7 h post ethanol treatment, left) and matured oocytes (7 h post progesterone treatment, right). (D) Tail-length dependent and independent translational regulation of CPE-containing mRNAs in frog oocytes. Shown for the CPE and a related motif at the top are translational efficiencies, as measured by the fold enrichment with pelleted ribosomes, of mRNAs in the N60-PAS*^mos^* library that contained the indicated number of motifs. Shown at the bottom are mean poly(A)-tail lengths of mRNAs in the N60-PAS*^mos^* library that contained the indicated number of motifs. Results are shown for prophase I-arrested oocytes (7 h post ethanol treatment, left) and matured oocytes (7 h post progesterone treatment, right); * indicates adjusted *P* value < 0.05, one-sided binomial test. (E) Translational repression by C-rich motifs and the CPE in frog oocytes. On the left are the schematics of reporters that examine the effects of motifs in two different 3ʹ-UTR sequence contexts. On the right are NanoLuc signals for each reporter after normalization to a co-injected firefly reporter. Error bars indicate standard deviation of biological triplicates; *P* values are from Student’s *t*-tests between indicated groups; WT, wild-type. (F) Translational and tail-length regulation by 3ʹ-UTR sequence motifs in frog embryos. This panel is as in (B), but for embryos at stage 6 (left) and stage 12 (right).

In frog oocytes undergoing maturation, we observed gradually increased TE for *k*-mers that contained the CPE (Figure 5B), a trajectory matching their tail-length changes (Figure 1H). Indeed, the TE associated with 6-mers strongly correlated with mean poly(A)-tail length in matured oocytes (*R*_p_ = 0.89, Figure 5C). Controls confirmed that these results depended on progesterone treatment and were sensitive to EDTA treatment that disrupted 80S ribosomes (Figure 5B–C, S5A). These results were consistent with the strong coupling between poly(A)-tail length and TE for endogenous mRNAs in frog oocytes.^26,41^ The high correlation also indicated that most translational control mediated by 3ʹ-UTR sequences occurs through effects on poly(A)-tail length.

A group of 6-mers containing multiple contiguous Cs did not follow the global trend and were associated with more translational repression without a corresponding decrease in tail length (Figure 5B–C). This repression was not maturation-specific; it was observed at all time points in both progesterone-treated oocytes and ethanol-treated control oocytes (Figure 5B). Stronger repression was observed when 3ʹ UTRs contained more C-rich *k*-mers, again without significant differences in tail lengths (Figure S5B). To rule out the possibility of non-translation-related depletion artifacts caused by the sucrose cushion, we measured TE with a different method, which was based on immunoprecipitating nascent chain-associated mRNAs that were actively translated (Figure S5C). mRNAs with C-rich *k*-mers were also depleted in this dataset (Figure S5D). These results indicated that translation of mRNAs containing the C-rich *k*-mers is repressed in frog oocytes in a manner that is independent of tail length.

Interestingly, although TE and poly(A)-tail length of injected mRNAs with CPE-containing 6-mers had almost a perfectly linear positive relationship in matured oocytes, in control oocytes not undergoing maturation, the TEs of these mRNAs were substantially lower than expected based on their longer-than-average poly(A) tails, implying that they were translationally repressed (Figure 5C). Although translational repression of CPE-containing mRNAs had been reported in prophase I-arrested oocytes,^56–58^ whether such repression depended on poly(A)-tail length had not been determined. Results from our mRNA library demonstrated that the CPE-mediated translational repression was tail-length independent. This repression also increased substantially when more CPEs were present within the 3ʹ UTR (Figure 5D), consistent with previous observations.^12,58^ Moreover, in prophase I-arrested oocytes, one CPE had strong synergistic translational repression with a second CPE but not with any other 5-mer (Figure S5E). The synergistic repression by multiple CPEs was highly specific because mRNAs that contained multiple copies of a CPE-like motif UUUUU were not repressed (Figure 5D, S5E).

To validate tail-length independent translational repression by the C-rich and CPE motifs, we measured luciferase activities of individual reporters injected into frog oocytes. We tested two 3ʹ-UTR sequences: one from the endogenous mRNA *krtcap2.L* and the other from a variant in the N60-PAS*^mos^* mRNA library, both of which contained two C-rich motifs (Figure 5E). The 3ʹ ends of the reporters were blocked with 3ʹ-dA to inhibit tail-length changes. In both sequence contexts, translation of the reporter containing C-rich motifs was repressed by more than 2-fold compared to that of reporters in which the C-rich motifs were mutated to non-repressing motifs (Figure 5E). When the non-repressing motifs were mutated to CPEs, translation of the reporters was also repressed, although to a lesser extent than that observed for the reporters with C-rich motifs.

An experiment examining the tail lengths and TEs of N60-PAS*^mos^* library injected into one-cell embryos yielded results for early embryos that resembled those observed for mature oocytes (Figure 5C and F), suggesting that early embryos inherit the polyadenylation and translational control regime operating in mature oocytes. The strong coupling observed between TE and poly(A)-tail length in oocytes and early embryos was substantially reduced, although not completely lost, by the time of gastrulation (22 hpi, stage 12), (Figure 5F), as expected from the switch of gene-regulatory regimes observed for endogenous mRNAs at this developmental period.^9^

Together, these results from our translational profiling of injected RNAs revealed an exceedingly tight relationship between CPE-dependent polyadenylation and enhanced translation during oocyte maturation and early embryonic development. Our results also showed that the CPE and C-rich motifs can cause tail-length-independent translational repression.

### Tail-length control during oocyte maturation is conserved among frogs, mice, and humans

To extend our analyses to mammals, we measured poly(A)-tail lengths of mouse mRNAs isolated from germinal vesicle (GV) oocytes and metaphase II (MII) oocytes that matured in vivo. As in frog oocytes, mRNAs in mouse oocytes underwent substantial tail-length changes during maturation^29,59^ (Figure S6A). Such changes strongly correlated with TE changes^29,59^ (Figure 6A), as observed for both fly^10,11^ and frog oocytes.^26^ As in frog oocytes, ROC analyses indicated that the UUUUA 5-mer had the greatest AUC value among all *k-*mers with lengths 3–8 nt (Figure S6B–C), and it out-performed the canonical CPEs or any combinations of previously reported CPEs (Figure 6B). Analysis of published TE and tail-length datasets from human oocytes^28,60^ yielded similar results (Figure 6C–D, Figure S6D–E). Moreover, the contextual features of the UUUUA element found in frog oocytes also appeared to function in both mouse and human oocytes (Figure 6E–F). Together, these analyses indicated that the UUUUA 5-mer specifies cytoplasmic polyadenylation in mouse and human oocytes, and the principles of poly(A) tail-length control are highly conserved among maturing oocytes of frogs, mice, and humans.

**Figure 6.**
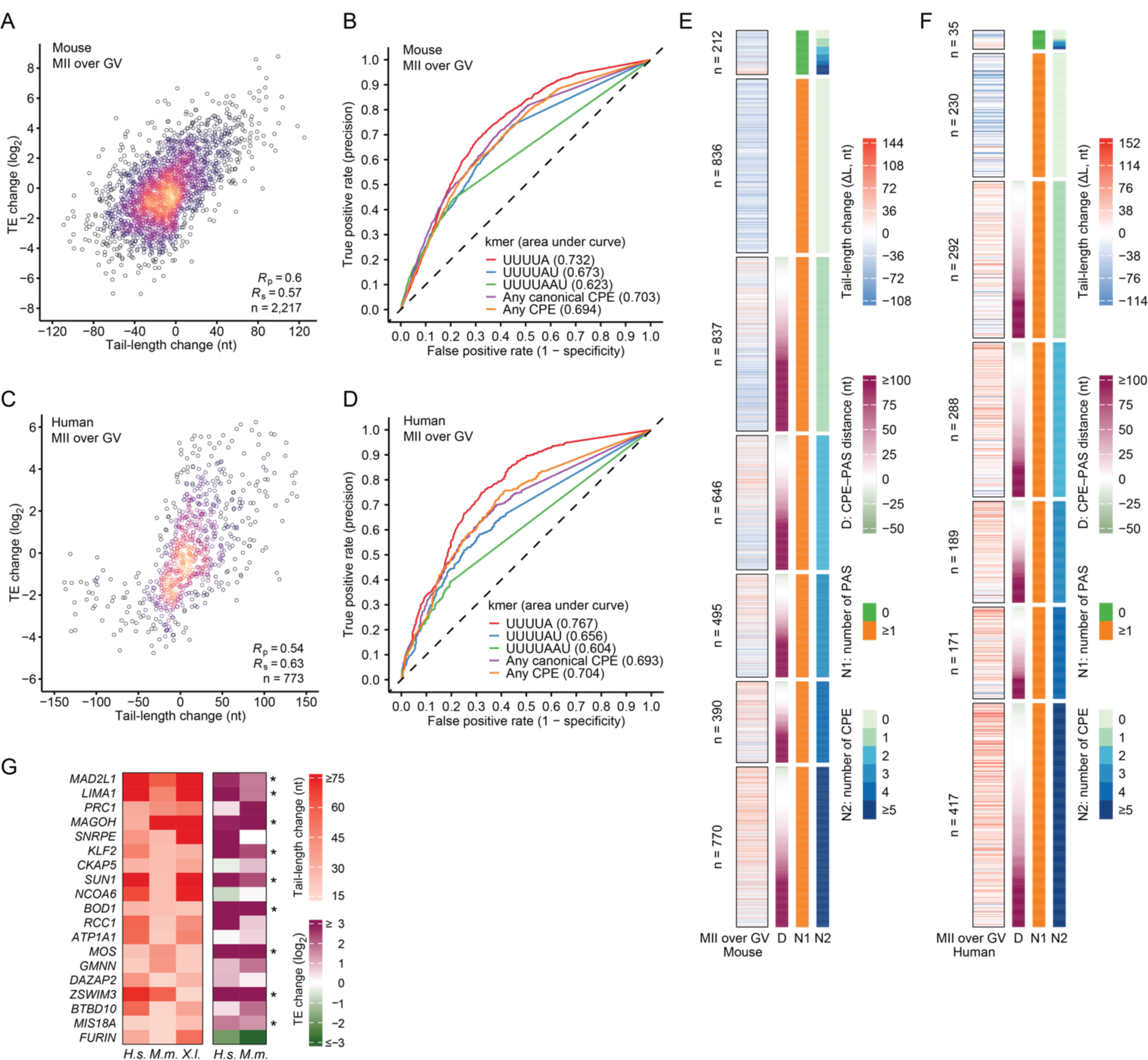
Tail-length control is conserved in mouse and human oocytes. (A) Coupling of TE with tail-length in mouse oocytes. Shown is the relationship between differences in TE and differences in poly(A)-tail length for mouse oocyte mRNAs, comparing values measured in GV oocytes to those measured in MII oocytes. For genes that contained multiple mRNA isoforms with different 3ʹ ends, only those that contained one dominant isoform (>90% by tag count) were included, and this dominant isoform was chosen to represent that gene. Colors indicate the density of points. (B) ROC curves testing the ability of the UUUUA and previously defined CPE motifs to classify endogenous mRNAs as subject to cytoplasmic polyadenylation during mouse oocyte maturation; otherwise, as in Figure 1F). (C) Coupling of TE and tail-length in human oocytes. As in (C), except differences were compared between values measured in human GV oocytes and those measured in human MII oocytes. (D) ROC curves testing the ability of the UUUUA and previously defined CPE motifs to classify endogenous mRNAs as subject to cytoplasmic polyadenylation during human oocyte maturation; otherwise, as in (B). (E) The effect of 3ʹ-UTR sequence features on tail-length changes of mRNAs as mouse GV oocytes mature to MII oocytes; otherwise, as in Figure 2I. (F) The effect of 3ʹ-UTR sequence features on tail-length changes of mRNAs as human GV oocytes mature to MII oocytes; otherwise as in (E). (G) Genes with substantial mRNA tail-lengthening (≥ 15 nt) in human (*H.s.*), mouse (*M.m.*), and frog (*X.l.*) oocytes. Heatmaps show measured tail-length changes (left) and TE changes (right), comparing between human GV and MII oocytes, mouse GV and MII oocytes, and frog oocytes 0 and 7 h post progesterone treatment. Asterisks indicate genes with ≥ 2-fold TE changes.

Finally, we asked if conserved tail lengthening of homologous genes among different species might shed light on their functional significance in oocyte maturation and early embryogenesis. By comparing tail-length changes in frog, mouse, and human oocytes, we identified 19 genes whose mRNA poly(A) tails underwent substantial lengthening (≥ 15 nt) in all three species (Figure 6G). Of these 19, nine had ≥ 2-fold TE increases in both mouse and human oocytes (Figure 6G). Among these genes were those that encode proteins with known functions in meiosis (*MAD2L1*, *SUN1*, *MOS*). Others were involved in cell cycle regulation (*BOD1*, *MIS18A*), cytoskeleton regulation (*LIMA1*), mRNA-splicing regulation (*MAGOH*), and transcriptional control (*KLF2*). We suggest that timely up-regulation of these proteins during oocyte maturation, presumably by tail-length-mediated translational activation, supports meiosis and early embryonic development.

## DISCUSSION

Although cytoplasmic polyadenylation during oocyte maturation has been studied for decades, the functional sequence motif responsible for poly(A)-tail lengthening, i.e. the CPE, had been only loosely defined, based on a handful of frog oocyte mRNAs. Analyzing tail-length sequencing of our diverse reporters, we identify a core 5-mer motif UUUUA that serves as the primary CPE in frog oocytes. This motif is shorter than anticipated, which explains why it had been previously missed in many mRNAs that undergo cytoplasmic polyadenylation. Yet despite its short length, the UUUUA CPE outperformed any combination of previously proposed longer CPEs in predicting tail lengthening for both injected and endogenous mRNAs. Moreover, this motif continued to drive the strongest polyadenylation in early frog embryos.

The UUUUA CPE element also specifies cytoplasmic polyadenylation in both mouse and human oocytes—with similar contextual features as in frog oocytes, thus underscoring the evolutionary conservation of the tail-length-mediated gene-regulatory mechanism under these developmental contexts. However, the CPE identified in zebrafish embryos differed from that of frog, mouse, and human, in that it permitted either A or U at the last position of the 5-mer (UUUUW). This increased degeneracy might have resulted from the binding of two alternative CPEB paralogs: CPEB1 (cpeb1a/b in fish) and CPEB4 (cpeb4a/b in fish). As indicated from in vitro binding assays^42,61^ and in vivo crosslinking sites,^43,62^ CPEB1 prefers to bind sites ending in A, whereas CPEB4 favors sites ending in U. Moreover, in fish embryos, translation of the *CPEB1* mRNA, as measured by ribosome-footprint profiling, is only ∼3-fold^9,41^ more than that of the *CPEB4* mRNA, whereas in frog oocytes and embryos the difference is 80-fold, implying that in fish embryos but not frog oocytes and embryos expression of CPEB4 protein might reach a level that can substantially impact cytoplasmic polyadenylation.

Identification of the actual CPE enabled key contextual features of this element to be identified. Likewise, we identified contextual features of the PAS that influence its tail-lengthening activity. For both elements, the contextual features appear conserved in fish, mouse, and human. Interestingly, for the CPE, the flanking nucleotides that favorably influence activity extend beyond the sequence-specific interactions observed in the structure of CPEB1 and its RNA ligand^63^, and the same is true for structures of human CPSF bound to the PAS.^64,65^ Perhaps the flanking nucleotides that influence activity facilitate either early recognition of these elements at a step preceding the final bound state or cooperative binding of both elements. With the identification of the CPE and the contextual features that substantially influence its activity and that of the PAS, we propose a simplified paradigm of cytoplasmic polyadenylation in early development, in which only two context-modulated elements explain why poly(A) tails of some mRNAs are extended much more than those of others.

Besides cytoplasmic polyadenylation, deadenylation also played a critical role in temporally sculpting poly(A)-tail lengths. This deadenylation occurred in two major waves. The first wave took place after GVBD and lasted until the first hour after fertilization. During this period, deadenylation activity was strong and independent of sequence motifs, such that not only mRNAs lacking either a CPE or a PAS but also those containing CPEs in non-optimal sequence contexts underwent net deadenylation. This enabled maturing oocytes to concentrate translational resources on mRNAs that had the longest tails, particularly those coding for proteins essential for meiosis and fertilization, a strategy seemingly also used in fish, mice, and humans.

The second wave of deadenylation occurred shortly after fertilization and continued through early embryonic development, during a period in which tail lengthening became more permissive. Unlike the first wave, tail-length shortening during this period was driven by specific-sequence motifs within mRNA 3ʹ UTRs. We identified several sequence motifs associated with stage-specific deadenylation, working successively in shortening poly(A) tails of different mRNAs during embryonic development. This mode of tail-length control would allow phased translational repression and clearance of different sets of maternal mRNAs and thus ensure a smooth transition to the zygotic transcriptome. Deadenylation in these two waves likely involved multiple deadenylases, as both PARN^66,67^ and CCR4-NOT^68^ have been implicated. Moreover, mutations of BTG4, an adaptor protein for CCR4-NOT have been associated with mouse^69,70^ and human infertility,^71,72^ supporting the significance of deadenylation during these developmental stages.

These sequence-specific and global effects all work together to produce the complex tapestry of tail length changes observed for endogenous mRNA as follows: The context-modulated CPE and PAS specify sequence-specific tail lengthening against a backdrop of global tail shortening in the maturing oocytes and more permissive tail lengthening in the embryo, sculpted by the subsequent action of additional sequence elements that specify waves of tail shortening.

Adding another layer of regulation, the CPE and C-rich motif mediated tail length-independent translational repression. In prophase I-arrested oocytes, the CPE binding of CPEB1 is proposed to recruit both the poly(A) polymerase Gld2 and the deadenylase PARN, with the stronger activity of PARN causing a net shortening of poly(A) tails of CPE-containing mRNAs.^73^ However, in our profiling of the N60-PAS*^mos^* library in prophase I-arrested oocytes, we did not observe preferential deadenylation of mRNAs that contained a CPE; on the contrary, their poly(A) tails were slightly longer than the average, indicating that repression of CPE-containing mRNAs does not require either short tails or the act of deadenylation. The mechanism of repression presumably instead involves another CPEB1-binding protein, Maskin, an eIF4E-interacting protein that displaces eIF4G to prevent efficient translation initiation.^74^

With respect to the C-rich motifs, C nucleotides in 3ʹ UTRs generally disfavored cytoplasmic polyadenylation, which could, in principle, lead to shorter poly(A) tails and lower TE of mRNAs that contained C-rich motifs. However, our results indicated that C-rich motifs could repress TE without relying on tail shortening. A similar motif within the 3ʹ UTR of LOX mRNA, which is CU-rich, mediates translational repression in erythroid cells, in which poly(C)-binding proteins HNRNPK and PCBP1 bind to the motif and inhibit 60S ribosomal subunit joining.^75,76^ Perhaps these or other poly(C)-binding proteins similarly recognize the C-rich motifs of oocyte mRNAs and mediate their translational repression. Such repression might reduce synthesis of proteins not required for meiosis progression and early embryogenesis. Interestingly, this repression persisted throughout oocyte maturation and early embryonic development, and was only alleviated during gastrulation. However, examination of published proteomic data^77^ did not uncover any known poly(C)-binding protein that exhibited corresponding changes in protein levels. Further investigation will be required to identify proteins involved in repressing mRNAs containing the C-rich motifs under these developmental contexts and to examine how this translational repression might contribute to oocyte maturation and early embryonic development.

## ACKNOWLEDGMENTS

We thank Thy Pham, Sean McGeary, and other current and former members of the Bartel lab for helpful discussions, A. Lokapally and H. Sive for experiments with frog embryos, D. Tomasello and O. Paugois for experiments with fish embryos, and the Whitehead Institute Genome Technology Core for sequencing. This work is supported by NIH grant GM118135. D.P.B. is an investigator of the Howard Hughes Medical Institute.

## AUTHOR CONTRIBUTIONS

K.X. and D.P.B. conceived the project, designed the study, and wrote the manuscript. J.L. isolated RNA from mouse oocytes. K.X. performed all the other experiments. K.X. analyzed all the data.

---

**Figure S1.**
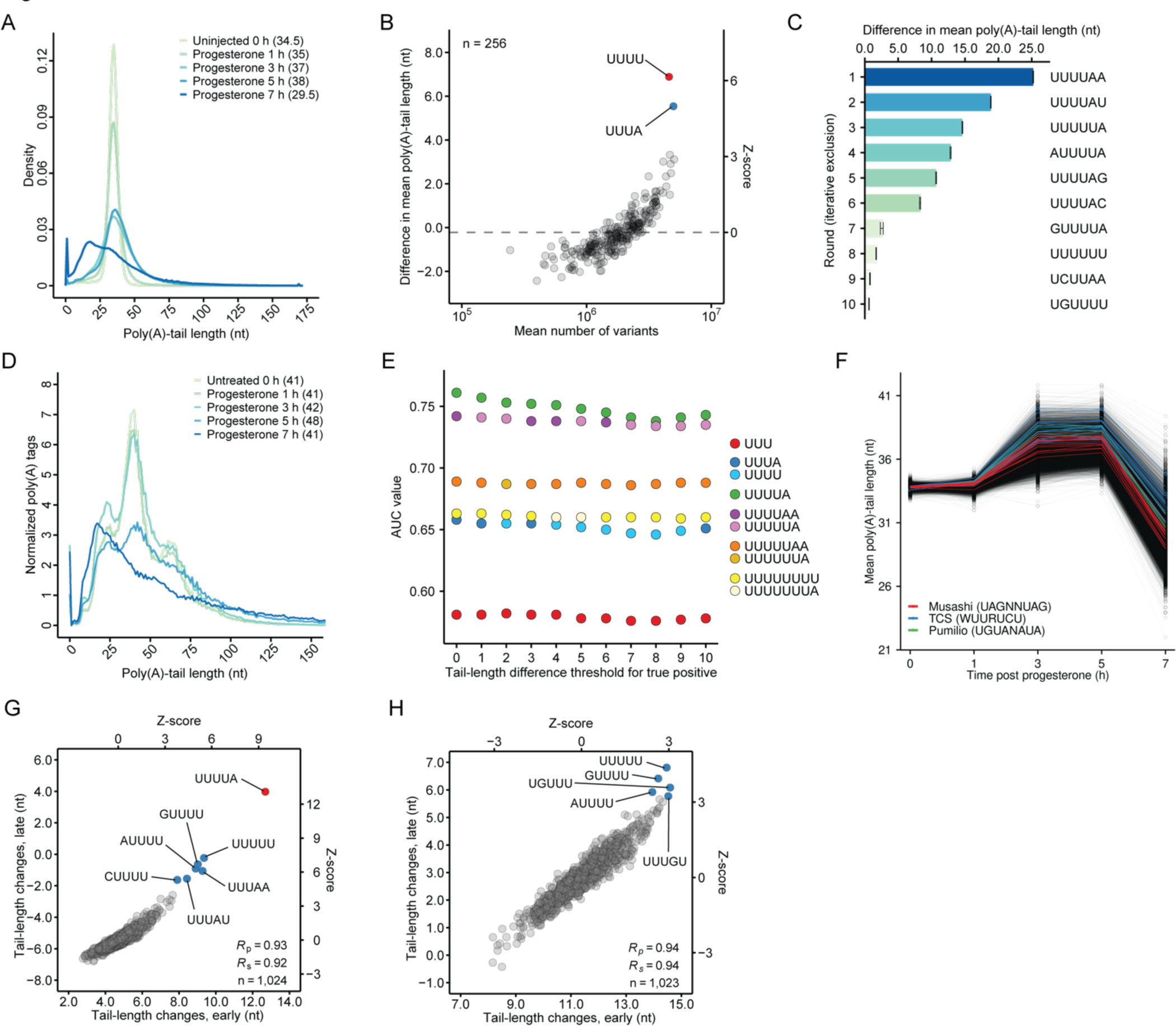
Supporting data regarding cytoplasmic polyadenylation in frog oocytes. Related to Figure 1. (A) Tail-length distributions of mRNAs of the N60-PAS*^mos^* library in oocytes collected at different times post progesterone treatment. Numbers in the parentheses indicate median tail lengths. (B) As in Figure 1C, but for 4-mers. (C) Top 6-mers associated with tail-length increases observed between 0 and 7 h post progesterone treatment. Tail-length differences were calculated in an iterative process with exclusion. In each round, tail-length differences were calculated for all 6-mers, and the 6-mer associated with the largest tail-length increase was selected. In the following rounds, all mRNA variants containing this 6-mer were excluded before this procedure was repeated. Note that in the 7^th^ round, the 6-mer GUUUUA can only be present in the last position of the variable region (a position at which the CPE has compromised activity; Figure 2A), because variants containing UUUUAA, UUUUAU, UUUUAG, and UUUUAC had all been excluded. (D) Tail-length distributions of endogenous mRNAs in frog oocytes collected at different times post progesterone treatment. Numbers in the parentheses indicate median tail lengths. (E) AUC values of ROC curves for best 3-, 4-, 5-, 6-, 7-, and 8-mer, when classifying cytoplasmic polyadenylation targets of endogenous mRNAs, as a function of the tail-length-difference threshold used in the classification. (F) No substantial polyadenylation associated with 8-mers lacking UUUUA. Plotted for each 8-mer are mean poly(A)-tail lengths for mRNAs containing that 8-mer, after excluding all variants of the N60-PAS*^mos^* library that contained UUUUA. Motifs previously implicated in cytoplasmic polyadenylation are colored. (G) Consistent tail-length changes associated with each 5-mer over time. Shown is the relationship between tail-length differences observed for mRNA variants in the N60-PAS*^mos^* library during late stages of oocyte maturation (7 h over 5 h) as a function of those observed earlier (3 h over 0 h). (H) No detectable maturation-specific cooperative effect observed between any 5-mer and the CPE. Plotted for each 5-mer, except the CPE, are differences in mean tail lengths for mRNAs containing that 5-mer, comparing late (7 h over 5 h) with early (3 h over 0 h) changes. Only variants that contained one UUUUA motif were included in this analysis.

**Figure S2.**
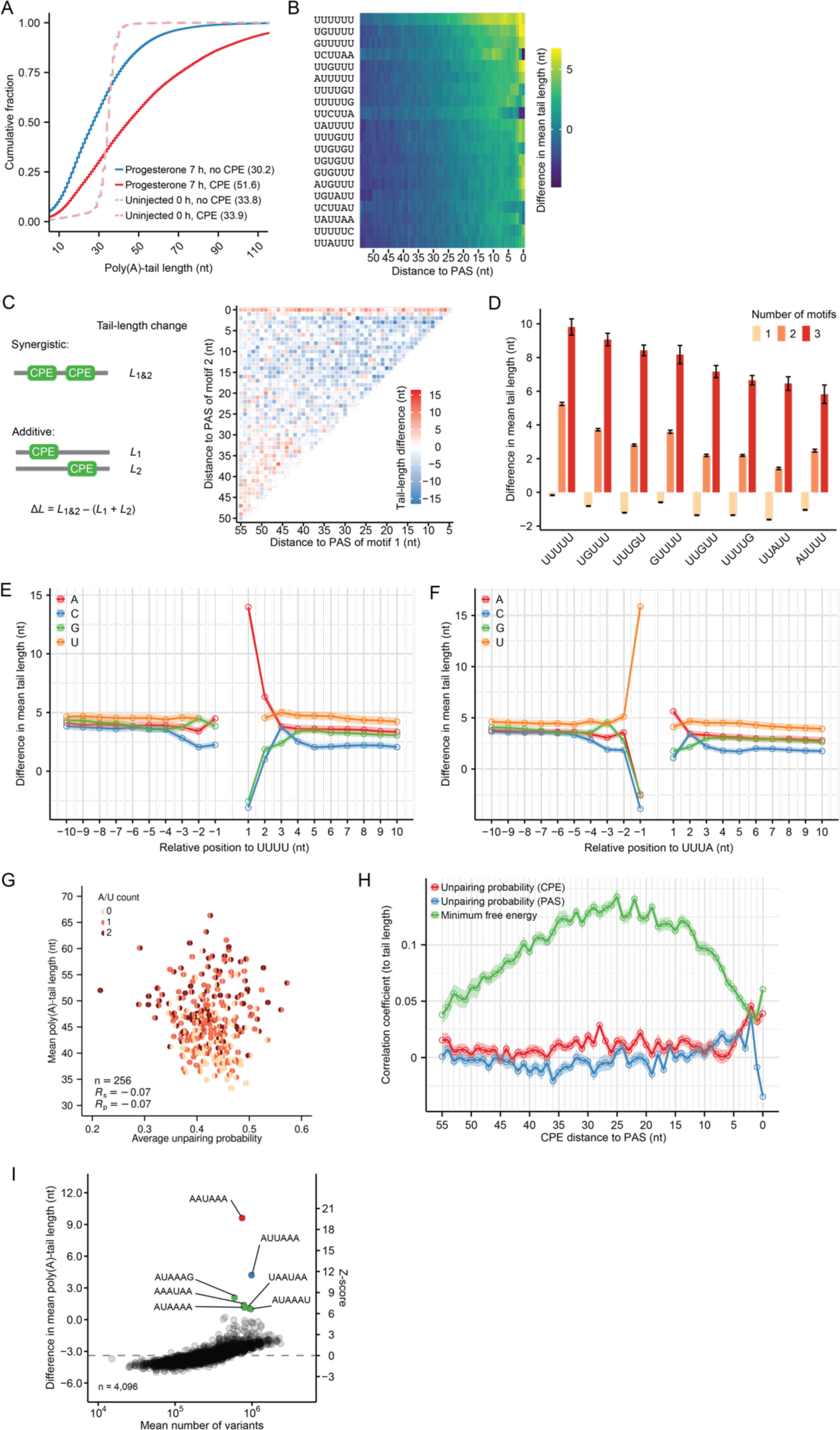
Supporting data regarding the ability of CPE and PAS contexts to modulate cytoplasmic polyadenylation in frog oocytes. Related to Figure 2. (A) Large variation of tail-length changes for CPE-containing mRNAs of the N60-PAS*^mos^* library. Shown are cumulative distributions of poly(A)-tail lengths for CPE-containing mRNAs and mRNAs that did not contain a CPE in oocytes at 0 and 7 h post progesterone treatment. Numbers in the parentheses indicate mean tail lengths. (B) Positional effects of CPE-like motifs. Heatmap shows the difference in mean poly(A)-tail length of mRNA variants in the N60-PAS*^mos^* library containing the indicated motifs at varying distances from the PAS, comparing between 0 and 7 h after progesterone treatment. Variants that contained a UUUUA CPE were excluded from this analysis. (C) Synergistic effects of two CPEs at some positions. The heatmap shows the difference between tail-length changes of variants that contained two CPEs and the sum of tail-length changes of variants that each contained a single CPE at the same locations. Tail lengths for mRNAs of the N60-PAS*^mos^* library were compared between 0 h and 7 h after progesterone treatment. (D) The influence of the number of CPE-like motifs on cytoplasmic polyadenylation. Shown is the difference in mean poly(A)-tail length of mRNA variants in the N60-PAS*^mos^* library containing different numbers of indicated CPE-like motifs but no CPE, comparing between 0 and 7 h after progesterone treatment. Error bars indicate standard error of the difference between means. (E) The effect of nucleotides flanking UUUU; otherwise as in Figure 2E. (F) The effect of nucleotides flanking UUUA; otherwise as in Figure 2E. (G) The relationship between mean poly(A)-tail length and average unpairing probability of the CPE in the N60-PAS*^mos^* library. Each circle represents a 9-mer containing a CPE flanked by 2 nt on each side. For each circle, the colors of the left and right semicircles indicate the A/U content of the 2 nt at the 5ʹ and the 3ʹ of the CPE, respectively. Only variants that contained a single CPE were included in this analysis. Average unpairing probability was predicted for the CPE by RNAfold.^78^ (H) Effects of predicted structural accessibility on poly(A)-tail lengths in the N60-PAS*^mos^* library. Plotted are average Pearson correlation coefficient (*R*p) values observed between structural accessibility scores predicted by RNAfold and mean poly(A)-tail lengths of variants that contained one CPE at varying distances to the PAS. For the CPE at each position, variants were divided into 20 equal-sized bins based on different GC contents in their random sequence-regions, and *R*p values were calculated for variants in each bin. Shaded areas indicate standard error. (I) Tail-length changes associated with each 6-mer in the 3ʹ UTRs of the CPE*^mos^*-N60 library, comparing between 0 and 5 h post progesterone treatment; otherwise as in Figure 1C.

**Figure S3.**
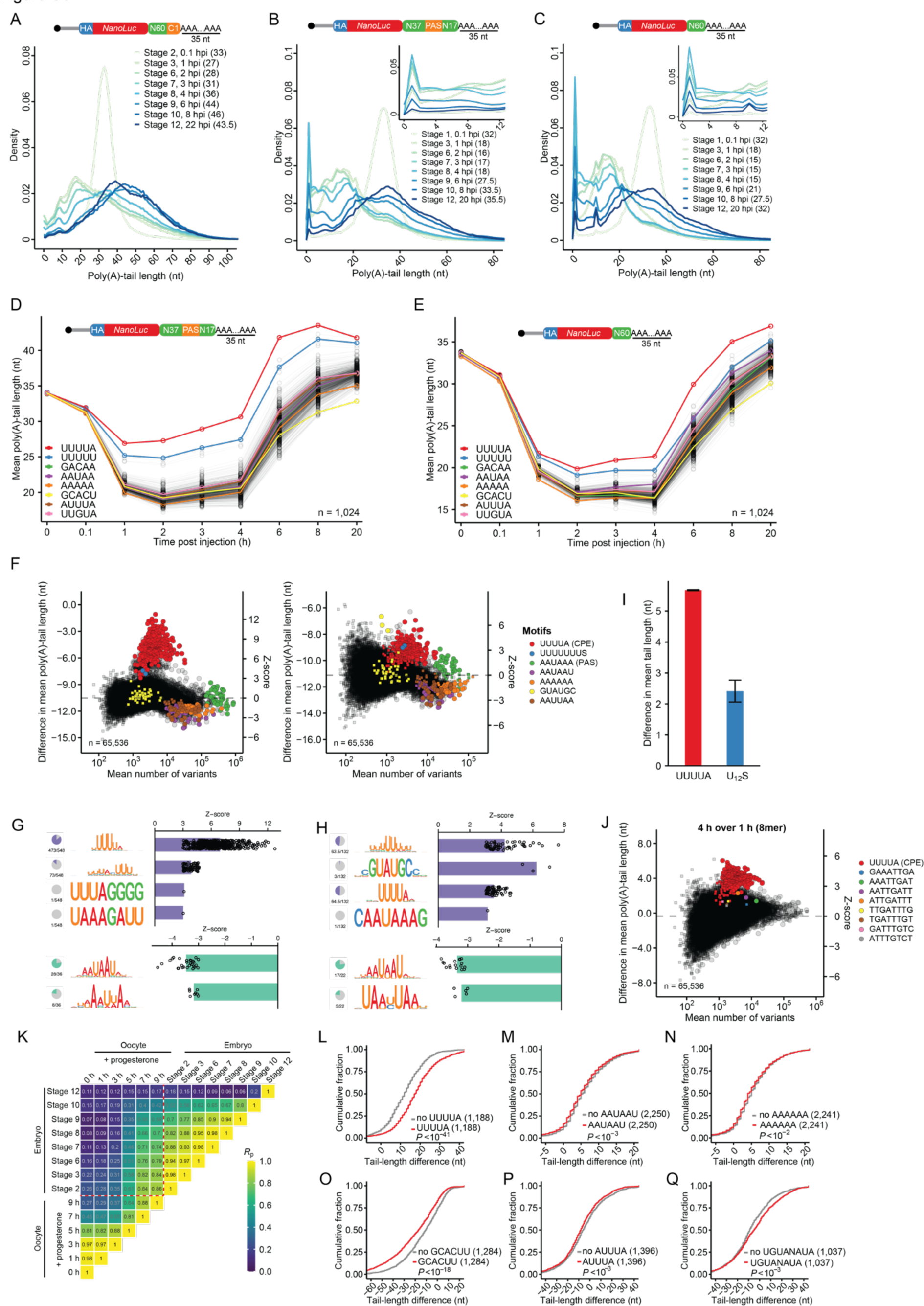
Supporting data for poly(A) tail-length control during frog early embryonic development. Related to Figure 3. (A) Tail-length distribution of members of the N60-PAS*^mos^* library (schematic as in Figure 1A) in frog embryos collected at different developmental stages. Numbers in the parentheses indicate median tail lengths. (B) As in (A), but for the N37-PAS-N17 library. The Inset shows a higher-resolution view of the short-tailed region. (C) As in (B), but for the N60 library. (D) As in Figure 3B, but for the N37-PAS-N17 library. (E) As in Figure 3B, but for the N60 library. (F) Tail-length changes associated with each 8-mer in the 3ʹ UTRs of mRNA variants in the N37-PAS-N17 library (left) or the N60 library (right), comparing between 0.1 h post injection (stage 2) and 1 h post injection (stage 3, S represents C or G); otherwise, as in the peripheral plots of Figure 3B. (G) Sequence motifs associated with tail-length changes during early frog embryogenesis. Shown are sequence logos generated from 8-mers most strongly associated with changes in tail length for members of the N37-PAS-N17 library, comparing between 0.1 h post injection (stage 2) to 1 h post injection (stage 3). The pie charts on the left indicate the fraction of 8-mers that aligned to the corresponding sequence logo. The bar plots on the right show the mean Z-scores, with points representing scores of individual 8-mers. (H) As in (G), but for the N60 library. (I) Effects of the CPE and U12 motifs on tail-length changes in frog embryos. Plotted are differences in mean poly(A)-tail lengths of mRNA variants in the N60-PAS*^mos^* library that had the indicated sequence motifs (S represents C or G), comparing between 0.1 h (stage 2) and 1 h (stage 3) post injection. Error bars indicate standard error of the difference between means. (J) Moderate but significant tail lengthening conferred by 8-mers residing within the *mos*.*L*-derived constant region of the N60-PAS*^mos^* library. This panel is as in (F), but for the N37-PAS-N17 library, comparing between 1 and 4 h post injection. In addition to coloring points for the 8-mers that contained the CPE, the points for eight different 8-mers tiled across the 15-nt sequence that separates the *mos*.L PAS from its poly(A) tail are also colored. (K) *R*p values reporting the tail-length correlation for each of the indicated pairwise comparisons. (L) Cumulative distributions of tail-length changes for endogenous frog mRNAs with 3ʹ UTRs that contained a CPE and those that did not, comparing between stage 2 and stage 8 frog embryos. The cohort that had more mRNAs was sub-sampled to match distributions of 3ʹ-UTR lengths and initial tail lengths of the other cohort. The numbers of mRNAs in each group are listed in parentheses; *P* value, Mann–Whitney U test. (M) As in L, but for the AAUAAU motif and comparing between stage 2 and stage 3 frog embryos. (N) As in L, but for the AAAAAA motif and comparing between stage 2 and stage 3 frog embryos. (O) As in L, but for the GCACUU motif (miR-427 target site) and comparing between stage 8 and stage 10 frog embryos. (P) As in L, but for the AUUUA motif (ARE) and comparing between stage 10 and stage 12 frog embryos. (Q) As in L, but for the UGUANAUA motif (PRE) and comparing between stage 10 and stage 12 frog embryos.

**Figure S4.**
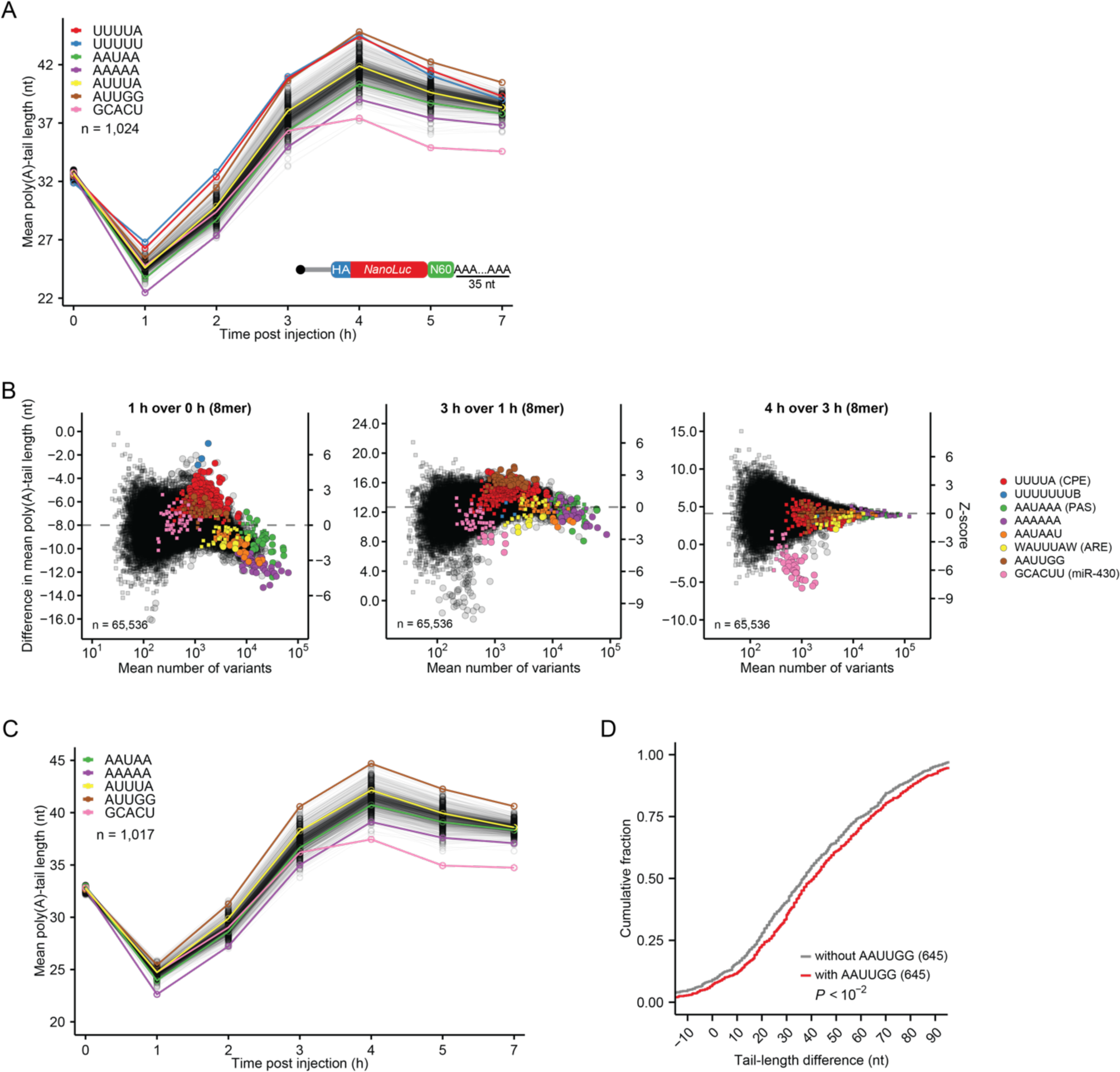
Supporting data for poly(A) tail-length control during zebrafish early embryonic development. Related to Figure 4 (A) As in Figure 4B, but for the N60 library. (B) As in Figure 4C, but for the N60 library. (C) As in panel A, except variants that contained either a UUUU motif or a PAS (AAUAAA or AUUAAA) were excluded from the analysis. (D) As in Figure 4E, but for the AAUUGG motif. The larger cohort (mRNAs without AAUUGG) was sub-sampled to match distributions of 3ʹ-UTR lengths and initial tail lengths.

**Figure S5.**
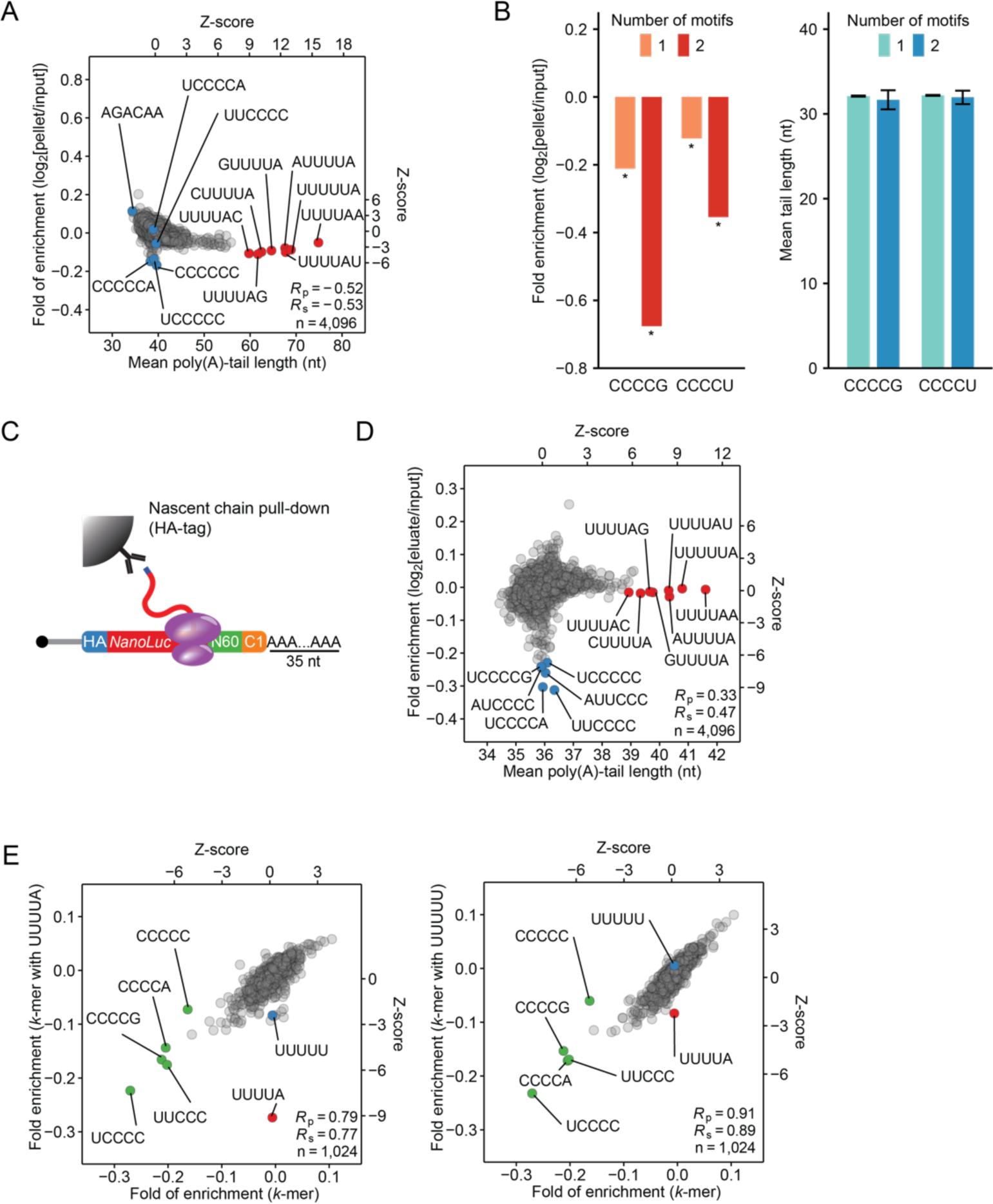
Additional data supporting tail-length-independent control of translation in frog oocytes. Related to Figure 5. (A) Evidence that translational profiling of injected mRNAs requires intact ribosomes. As in Figure 5C, except lysates were prepared in the presence of EDTA. (B) Tail-length-independent translational repression associated with C-rich motifs in frog oocytes. Shown are translational efficiencies (left) and poly(A)-tail lengths (right) of mRNAs containing indicated C-rich motifs in the N60-PAS*^mos^* library in prophase I-arrested oocytes (7 h post ethanol treatment); * indicates adjusted *P* < 0.05, one-sided binomial test. (C) Schematic of immunoprecipitation of mRNAs associated with nascent polypeptides. (D) Translational regulation by 3ʹ-UTR sequence motifs in prophase I-arrested frog oocytes, profiled by immunoprecipitation of mRNAs associated with nascent polypeptides. Shown for each 6-mer in the 3ʹ UTRs of the N60-PAS*^mos^* library is the mean translational efficiency (as measured by the fold enrichment in the immunoprecipitation) observed for mRNAs with that 6-mer, plotted as a function of mean poly(A)-tail length of these mRNAs. Results are shown for prophase I-arrested oocytes (7 h post ethanol treatment). Otherwise, this panel is as in Figure 5C. (E) Synergistic repression by two CPEs. Shown for each 5-mer in the 3ʹ UTRs of the N60-PAS*^mos^* library is its contribution to translational efficiency when it was co-present with either a CPE (left) or a CPE-like motif (UUUUU, right) within the same 3ʹ UTR, plotted as a function of its overall contribution to translational efficiency. Results are shown for prophase I-arrested oocytes (7 h post ethanol treatment). Translational profiling was performed with ribosome pelleting in a sucrose cushion.

**Figure S6.**
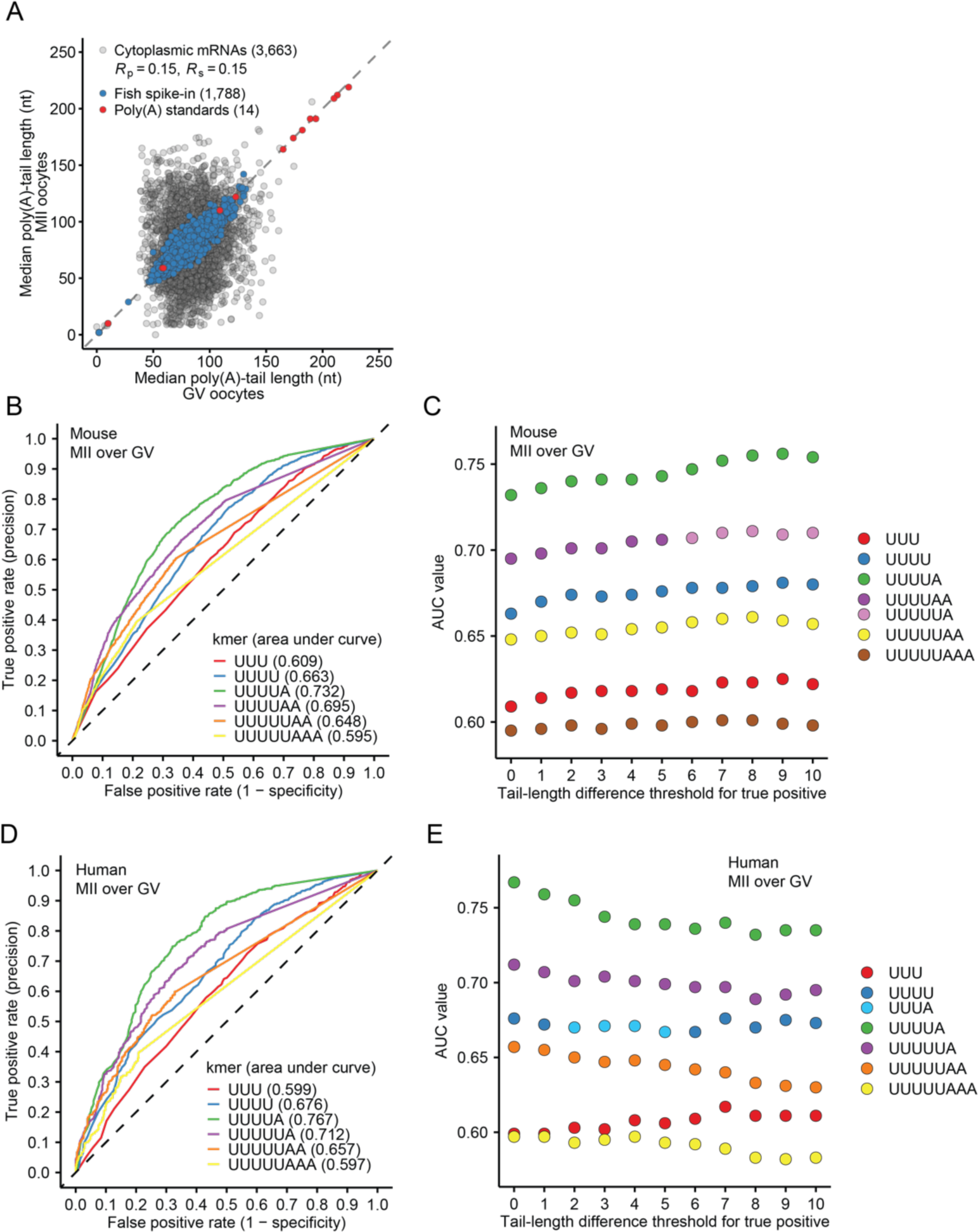
Additional data supporting conserved tail-length control in mouse and human oocytes. Related to Figure 6. (A) Poly(A) tail-length change during mouse oocyte maturation. Shown is the plot comparing median poly(A)-tail lengths in mouse MII oocytes to those in mouse GV oocytes. Results are shown for mouse cytoplasmic mRNAs (gray), zebrafish spike-in mRNAs, and poly(A) standards (red). Only RNAs with ≥50 tags were shown. Points for two poly(A) standards are outside the plotting area. (B) ROC curves testing the ability of the best 3-, 4-, 5-, 6-, 7-, and 8-mer to classify endogenous mRNAs as subject to cytoplasmic polyadenylation during mouse oocyte maturation. (C) AUC values of ROC curves for best 3-, 4-, 5-, 6-, 7-, and 8-mer, when classifying cytoplasmic polyadenylation targets of mouse endogenous mRNAs, as a function of the tail-length-difference threshold used in the classification. (D) ROC curves testing the ability of the best 3-, 4-, 5-, 6-, 7-, and 8-mer to classify endogenous mRNAs as subject to cytoplasmic polyadenylation during human oocyte maturation. (E) AUC values of ROC curves for best 3-, 4-, 5-, 6-, 7-, and 8-mer, when classifying cytoplasmic polyadenylation targets of human endogenous mRNAs, as a function of the tail-length-difference threshold used in the classification.

## METHODS

### Oligo sequences

All oligo sequences used in this study are listed in Table S1.

### Construction of DNA templates for in vitro transcription

For the N60-PAS*^mos^* library, the DNA template was assembled from four DNA fragments by overlapping PCR. Three out of four fragments were amplified using the KAPA HiFi HotStart Kit (Roche, KK2502) following the manufacturer’s suggested protocol, from the plasmid C071 with the following primer sets: KXU024 and KXR303 (for the T7 promoter and the 5ʹ-UTR, fragment 1 [F1]), KXF303 and KXU236 (for the *NanoLuc* coding region, fragment 2 [F2]), KXU110 and KXU068 (for the HDV ribozyme, fragment 4 [F4]). The other fragment (F3), which contained a 60-nt random-sequence region (A:C:G:T = 30:19:19:32), a 21-nt region based on the 3ʹ-end of frog *mos.L* mRNA, and a 35-nt poly(A) stretch was constructed by Klenow extension of oligos KXS050 and KXS051. The two oligos (200 pmol each) were annealed in water in a 49-µl reaction, which was incubated at 85°C for 5 min and then slowly cooled to 25°C at 0.1°C/sec. A mixture containing 6 µl NEBuffer 2 (New England Biolabs, B7002S), 2 µl Klenow enzyme (5 units/µl, New England Biolabs, M0210S), and 3 µl 10 mM dNTP (KAPA HiFi Kits from Roche, KK2502) were added to the annealed oligos. The reaction was incubated at 25°C for 15 min and stopped by adding 1.2 µl 500 mM EDTA and incubating at 75°C for 20 min. Overlapping PCR reactions were performed to join F1 and F2, and F3 and F4 separately. The joined products were then joined to form the final DNA product. A standard overlapping PCR was performed in a 100-µl reaction containing 10 pmol of each fragment, 2 µl KAPA HiFi enzyme, 3 µl 10 mM dNTP, and 20 µl 5x KAPA HiFi buffer for 15 cycles, with an annealing temperature of 66°C. All PCR products and double-stranded DNAs were purified with agarose gels (Lonza, 50004) and the GeneJet Gel Extraction Kit (Thermo Fisher, K0692).

DNA templates for the N60 and the N37-PAS-N17 libraries were constructed similarly except for F3. For this fragment, a splint ligation was carried out with an acceptor oligo containing the variable region(s) (KXS088 for the N60 library and KXS087 for the N37-PAS-N17 library), a donor oligo KXS053, and the splint oligo KXS055. The oligos (1 nmol each) were mixed in water in a 50-µl reaction, heated to 95°C for 5 min, and then slowly cooled to 25°C at 0.1°C/sec. 24 µl was taken from the annealed product, mixed with 3 µl T4 DNA ligase and 3 µl T4 DNA ligase buffer (New England Biolabs, M0202S), and incubated at 25°C overnight (16 hr). The ligated product was mixed with an equal volume of Gel Loading Buffer II (Thermo Fisher, AM8547), heated for 5 min at 95°C, and resolved on an 8% urea-acrylamide denaturing gel (National Diagnostics, EC-828). The band migrating at the size of 138 nt was identified by UV-shadowing, excised, macerated, and eluted in a buffer containing 10 mM HPEPS pH 7.5 and 300 mM NaCl at 70°C for 30 min. The gel pieces were removed using Spin-X columns (Corning, 8160). DNAs were then precipitated with isopropanol and resuspended in water for downstream Klenow extension with oligo KXU237 in the same way as F3 in the N60-PAS*^mos^* library.

The DNA template for the CPE*^mos^*-N60 library was constructed in a similar way to that for the N60 library, except for the following differences: 1) F1 and F2 were amplified as one fragment from the plasmid C071 with primers KXU024 and KXS054 (thus this library lacks the HA tag); 2) KXS052 was used as the acceptor oligo for the splint ligation when F3 was generated.

For individual mRNA reporters used for luciferase assays, DNA templates were assembled with fragments F1 and F2 as in the DNA template for the N60-PAS*^mos^* library, except that F2 was amplified with a different reverse primer (one of KXU333–KXU338). Each reverse primer contained a different 3ʹ-UTR sequences and a 35-nt poly(A) tail.

### Preparation of mRNAs for injections

For mRNA libraries, in vitro transcription was performed in a 100-µl reaction containing 40 mM Tris pH 8.0, 21 mM MgCl_2_, 2 mM Spermidine (Sigma, 85558-1G), 1 mM dithiothreitol (GoldBio, DTT25), 5 mM NTP Set (Thermo Fisher, R0481), 0.2 units yeast inorganic pyrophosphatase (NEB), 80 units SUPERase•In (Thermo Fisher, AM2694), 2 µg DNA template, and T7 RNA Polymerase (purified in-house and used at final concentration of 6.4 ng/µl). After incubation at 37°C for 3 hr, 2 units of DNase I (New England Biolabs, M0303S) were added, followed by another 20 min incubation at 37°C. To enhance HDV ribozyme cleavage, thermal cycling was performed (65°C for 90 sec and 37°C for 5 min, four cycles, 50 µl of reaction per tube). Before gel loading, 2 µl 0.5 M EDTA pH 8.0 and 100 µl Gel Loading Buffer II were added. After incubation at 65°C for 5 min, RNAs were separated on 5% urea-acrylamide denaturing gels. Desired RNA bands were identified by UV-shadowing, excised, macerated, and eluted in a buffer containing 10 mM HPEPS pH 7.5 and 300 mM NaCl at 23°C overnight (> 16 hr) on a thermomixer, shaking at 1400 rpm (15 sec on and 105 sec off). The gel pieces were removed using Spin-X columns (Corning 8160), and RNAs were precipitated with isopropanol and resuspended in water for downstream reactions.

Capping of RNAs was carried out with the Vaccinia Capping System (New England Biolabs, M2080S) following the manufacturer’s protocol, except that the Vaccinia capping enzyme was used at 2 units/µl. RNAs were then purified by phenol/chloroform extraction and ethanol precipitation. Water-resuspended RNAs were applied to Micro Bio-Spin P-30 columns (Bio-Rad, 7326250) for desalting.

3ʹ-end cyclic phosphates generated by HDV ribozyme cleavage were removed in a 100-µl reaction containing up to 100 µg capped RNAs, 50 units T4 Polynucleotide Kinase (PNK, New England Biolabs, M0201S), 1x T4 PNK buffer, and 25 units SUPERase•In. After incubation at 37°C for 1 hr, RNAs were purified by phenol/chloroform extraction and ethanol precipitation. RNAs were then resuspended in 1x Gel Loading Buffer II and purified with urea-acrylamide gels in the same way as after in vitro transcription. All RNAs were checked for integrity by visualization on formaldehyde-agarose denaturing gels as described^41^ before being stored at –80°C.

Individual mRNAs used for luciferase assays were in vitro transcribed the same way as mRNA libraries but in 50-µl reactions, and no HDV cycling was performed after the DNase-I digestion. RNAs were supplemented with 2 µl 0.5 M EDTA pH 8.0 and applied to Micro Bio-Spin P-30 columns for desalting. RNAs were then purified by phenol/chloroform extraction and ethanol precipitation. Capping was performed the same as for the mRNA libraries. 3ʹ-dATP was added to RNAs with Yeast Poly(A) Polymerase (ThermoFisher, 74225Z25KU) in a 40-µl reaction containing 1x Yeast Poly(A) Polymerase Reaction Buffer, 4 µM RNA, 500 µM 3ʹ-dATP (Jena Bioscience, NU-1123L), 1200 units Yeast Poly(A) Polymerase, and 20 units SUPERase•In. After incubation at 37°C for 1 h, RNAs were applied to Micro Bio-Spin P-30 columns, and the flowthrough was further purified by phenol/chloroform extraction and ethanol precipitation. To confirm that the 3ʹ-ends were blocked by 3ʹ-dATP, poly(A) tailing assays were performed with purified RNAs in a 10-µl reaction containing 1x Yeast Poly(A) Polymerase Reaction Buffer, 500 ng RNA, 500 µM ATP (New England Biolabs), and 300 units Yeast Poly(A) Polymerase. After 20 min incubation at 37°C, RNAs were resolved on formaldehyde-agarose denaturing gels as described.^41^

### Mouse husbandry

The C57BL/6J inbred mouse line was used in this study. Mice were housed in a 12-12 h light/dark cycle with constant temperature and food at the Whitehead Institute for Biomedical Research. The Massachusetts Institute of Technology Department of Comparative Medicine provided daily cage maintenance and veterinarian health checks. All animal experiments performed in this study were approved by the Massachusetts Institute of Technology Committee on Animal Care.

### Frog oocytes and embryos and fish embryos

Frog oocytes used for the injection of the N60-PAS*^mos^* and the CPE*^mos^*-R60 libraries and measurements of endogenous mRNA poly(A)-tail lengths were obtained from frog (*X. laevis*) ovaries purchased from Nasco (LM00935) as described.^41^ Due to the discontinuation of frog ovaries from Nasco, frog oocytes used for injections of the N60, N37-PAS-N17, and CPE*^mos^*-N60 libraries as well as individual mRNA reporters were obtained from Xenopus1 (12005). Because these oocytes were defolliculated by the vendor, healthy Stage V and VI oocytes were hand-picked and transferred to OR-2 buffer (5 mM HEPES pH 7.6, 82.5 mM NaCl, 2.5 mM KCl, 1 mM MgCl_2_, 1 mM CaCl_2_, 1 mM Na_2_HPO4) supplemented with 100 µg/ml Gentamicin (Thermo Fisher, 15750060) incubated at 18°C overnight (> 16 h) for recovery before injections.

Frog oocytes were matured in vitro in OR-2 buffer supplemented with 10 µg/ml progesterone (Millipore Sigma, P0130) from a 10 mg/ml stock in ethanol. For controls, oocytes were incubated in OR-2 buffer with ethanol (0.1%). The time required for frog oocytes to mature varied between different batches. In most cases, 50% of oocytes reached GVBD (displaying a white spot in the animal pole) between 3 and 5 h, and 100% of oocytes reached GVBD by 7 h after incubation with progesterone.

For frog embryos, female frogs (*X. laevis*) were injected with 700 units of human chorionic gonadotropin (hCG) into the dorsal lymph sac. After 12–16 h, eggs were harvested and fertilized in vitro with testis pieces cut out from a sacrificed male frog in 0.1x MBS (8.8 mM NaCl, 0.1 mM KCl, 0.1 mM MgSO_4_, 0.5 mM HEPES, 0.25 mM NaHCO_3_, 0.07 mM CaCl_2_, pH 7.8). The fertilized eggs were de-jellied in 2% (w/v) cysteine (pH 8.0), thoroughly rinsed with 0.1x MBS, and transferred to 1x MBS for injections. After injections, frog embryos were kept in 1x MBS at 23°C until stage 10, when they were moved to 15°C until they reached stage 12. Frog embryos were staged as described.^79^

Zebrafish (*Danio rerio*) embryos were obtained from the wide-type AB line, incubated in E3 medium (5 mM NaCl, 0.17 mM KCl, 0.33 mM CaCl_2_, 0.33 mM MgSO_4_, 0.1% w/v Methylene Blue) at 28°C, and staged as described.^80^

All animal use was in accordance with a protocol approved by the Massachusetts Institute of Technology Committee on Animal Care.

### Oocyte and embryo injections

All injections were performed at 23°C with a PLI-100 Plus Pico-Injector. Stage V or VI frog oocytes that were healthy after overnight recovery were selected for injection. For the N60-PAS*^mos^* and CPE*^mos^*-N60 libraries, 4 nl mRNA (0.5 pmol/µl) was injected per oocyte. The N60 and N37-PAS-N17 libraries were mixed at a 1:3 molar ratio, and 4 nl (0.5 pmol/µl) of the mix was injected per oocyte. For luciferase assays, each oocyte was co-injected with a NanoLuc luciferase reporter mRNA with a unique 3ʹ-UTR sequence (0.05 pmol/µl) and a firefly luciferase mRNA with 120-nt poly(A) region followed by a mutant mouse Malat1 3ʹ-end (0.1 pmol/µl), which was used for normalization.^41^

Frog embryos were injected at stage 1. For the N60-PAS*^mos^* library, 4 nl mRNA (0.5 pmol/µl) was injected per embryo. The N60 and N37-PAS-N17 libraries were mixed at a 1:3 molar ratio, and 4 nl (0.1 pmol/µl) was injected per embryo. Fish embryos were injected at the 1-cell stage. The N60 and N37-PAS-N17 libraries were mixed at a 1:3 molar ratio in a buffer containing 0.025% phenol red and 100 mM KCl. Each embryo was injected with 2 nl of the RNA library mix (0.1 pmol/µl).

### Sample collection for poly(A) tail-length and translational profiling

Frog oocytes (50–100) were collected at the indicated time after either ethanol or progesterone treatment. OR-2 buffer was removed and oocytes were washed three times with ice-cold buffer RL (20 mM HEPES pH 7.5, 100 mM KCl, 5 mM MgCl_2_, 1% [v/v] Triton X-100, 100 µg/ml cycloheximide, cOmplete protease inhibitor cocktail [MilliporeSigma, 11836170001, 1 tablet per 10 ml buffer], 10 µl/oocyte). After removing all wash buffer, oocytes were lysed in buffer RL supplemented with 200 units/ml SUPERase•In (10 µl/oocyte) by vigorous shaking and pipetting. Lysates were cleared by centrifugation at 5000*g* at 4°C for 10 min. A portion of each supernatant (50 µl, equivalent to 5 oocytes) was mixed with 500 µl Tri Reagent (Thermo Fisher, AM9738). The rest of each supernatant was either used for nascent-chain pulldown immediately or flash-frozen in liquid nitrogen in aliquots stored at –80°C for analysis on a sucrose cushion. Note that for reporter mRNAs, the “0 h post progesterone treatment” sample was prepared by collecting the supernatant of the lysate of untreated oocytes and combining it with the uninjected mRNA library before the Tri Reagent was added to the mix. For endogenous mRNAs, the “0 h post progesterone treatment” sample was prepared by collecting the supernatant of the lysate of oocytes not treated with any reagents.

Frog embryos (∼30) were collected at indicated developmental stages the same way as frog oocytes. Fish embryos (10–15) were collected at indicated developmental stages, E3 medium was removed, and embryos were washed once with 1 ml E3 medium and incubated in 1 ml E3 medium with 2 mg/ml pronase for 4 min for de-chorionation. After removing all E3 medium, 1 ml Tri Reagent was added to the embryos, followed by vigorous shaking and pipetting.

All RNA isolations with the Tri Reagent were performed with Phasemaker tubes (Thermo Fisher, A33248) according to the manufacturer’s protocol.

### Mouse oocyte and egg collection

For the collection of prophase I-arrested (GV) oocytes, 5- to 8-week-old female mice were superovulated by intraperitoneal injection of 5–10 I.U. of pregnant mare’s serum gonadotropin (PMSG) (Thermo Fisher, NC1663485). At 48 h post injection, the mice were euthanized, and the ovaries were dissected into MEM (Sigma Aldrich, M0268), containing 25 mM HEPES pH 7.3, 0.1 mg/mL sodium pyruvate (Sigma Aldrich, P4562), 3 mg/mL polyvinylpyrrolidone (Sigma Aldrich, P2307), 10 μg/mL gentamicin (Life Technologies, 15710064), and 2.5 μM milrinone (Sigma Aldrich, M4659). Oocytes were collected in 37°C MEM in the presence of milrinone to prevent maturation. Cumulus cells were removed from cumulus-oocyte-complexes by repeated aspiration through a glass pipette. GV oocytes were collected in TRI reagent, and total RNA was isolated according to manufacturer’s instructions. Approximately 100 oocytes from multiple animals were collected per biological replicate.

For the collection of metaphase II (MII) oocytes, 5- to 8-week-old female mice were injected with 5–10 I.U. of PMSG. At 48 h after PMSG injection, the animals were injected with 5–10 I.U. of Human chorionic gonadotropin (hCG) (Millipore Sigma, CG5-1VL). At 16 h post hCG injection, cumulous-enclosed egg complexes were collected from the oviduct and cultured in 37°C MEM with hyaluronidase (3 mg/ml, Sigma Aldrich, H4272) for 2 min. Denuded eggs were then washed free of hyaluronidase, collected in TRI Reagent, and RNA was isolated according to manufacturer’s instructions. Approximately 100 MII oocytes from multiple animals were collected per biological replicate.

### Sucrose cushion for translational profiling

Frog oocyte or embryo lysate (∼200 µl) was laid on top of an ice-cold 1.8-ml sucrose cushion (20 mM HEPES pH 7.5, 100 mM KCl, 5 mM MgCl_2_, 1% [v/v] Triton X-100, 1 M sucrose, 100 µg/ml cycloheximide, 20 units/ml SUPERase•In) in a polycarbonate tube (Bechman Coulter, 362305). Ribosomes were pelleted by centrifugation at 110,000 rpm (657,000*g*) at 4°C in a Beckman TLA-100 rotor for 1 h. After centrifugation, the supernatant was removed, and 1 ml Tri Reagent was added to the pellet.

### Nascent-chain pulldown of ribosome-associated RNAs

Frog oocyte or embryo lysate was incubated with anti-HA magnetic beads (Thermo Fisher, 88837) using 20 µl slurry per 500 µl lysate. The bead mixture was incubated at 4°C for 1 h with end-to-end rotation. Beads were then immobilized using a magnetic stand, and the supernatant was removed. Beads were washed three times with 0.6 ml buffer RL, resuspended in 100 µl buffer RL, mixed with 1 ml Tri Reagent and saved for RNA isolation.

### Luciferase assays

Oocyte lysates for luciferase assays were prepared as described.^41^ Luciferase assays were performed with Nano-Glo Dual-Luciferase Reporter Assay System (Promega, N1630) in a GloMax Discover plate reader according to the manufacturer’s protocol.

### 3ʹ-UTR variant and tail-length sequencing

In some cases, total RNA containing reporter mRNA libraries was ligated to a pre-adenylated 3ʹ adapter directly, but for the N60 library injected into fish embryos and frog oocytes, the N37-PAS-N17 library injected into fish embryos and frog oocytes, and the CPE*^mos^*-N60 library injected into frog oocytes, reporter library mRNAs were first enriched by hybridization with biotinylated oligos. RNA isolated from the oocyte or embryo lysate was mixed with 8 pmol KXSH009, 8 pmol KXSH010, and 2x SSC (0.3 M NaCl, 30 mM sodium citrate pH 7.0) in a total of 50 µl. The RNA and oligos were annealed by incubation at 70°C for 5 min and then slowly cooling to 23°C at 0.1°C/sec. The annealed mixture was combined with 40 µl MyOne Streptavidin C1 beads (Thermo Fisher, 65002) and incubated for 20 min at 23°C on a thermal mixer, shaking with 15 sec on and 1 min 45 sec off. The supernatant was separated from the beads with a magnetic rack and removed. The beads were washed twice with 300 µl 1xB&W buffer (5 mM Tris-HCl pH 7.5, 0.5 mM EDTA, 1 M NaCl) and once with 300 µl 2x SSC. The RNA was eluted from the beads first with 100 µl 10 mM HEPES pH 7.5 at 65°C for 3 min and then with 100 µl water at 65°C for 3 min. The eluates were combined, precipitated with ethanol, and resuspended in 6.5 µl water.

The enriched or unenriched RNA isolated from oocyte or embryo lysates was ligated to a pre-adenylated 3ʹ adapter in a 10 µl reaction containing 5 µM 3ʹ adapter KXS330, 50 mM HEPES pH 7.5, 10 mM MgCl_2_, 10 mM dithiothreitol, 1 unit/µl T4 RNA ligase 1 (New England Biolabs, M0204S) and incubated at 23°C for 150 min. After ligation, RNA was extracted with phenol/chloroform, precipitated with ethanol, and resuspended in 11.4 µl water. The ligated RNA was mixed with 0.6 µl 100 µM reverse transcription primer KXS037 in a total volume of 12 µl, incubated at 65°C for 5 min, and cooled on ice for 1 min. The annealed RNA was reverse transcribed in a 20 µl reaction containing 1x First-Strand Buffer, 500 µM dNTPs, 5 mM dithiothreitol, 1 unit/µl SUPERase•In, and 200 units SuperScript III (Thermo Fisher, 18080044) incubated at 50°C for 1 h. RNA was then hydrolyzed with addition of 3.3 µl 1 M NaOH and incubation at 90°C for 10 min, followed by neutralization with 36.7 µl 1 M HEPES pH 7.5, and the cDNA was collected by desalting with a Micro Bio-Spin P-30 column. The cDNA library was amplified in a 50 µl PCR reaction with KXS037 and a barcoded primer (KXS057) that hybridized to a constant region of the library, using the KAPA HiFi HotStart Kit following the manufacturer’s suggested protocol for 10–15 cycles. The PCR-amplified library was cleaned up twice with AMPure XP beads (Beckman Coulter, A63881) with a beads-to-sample ratio of 1.2.

Sequencing was performed on an Illumina HiSeq 2500, with a standard single-read run of 300 cycles and sequencing primer KXS067 (for the N60-PAS*^mos^*, N60, and N37-PAS-N17 libraries) or KXS056 (for the CPE*^mos^*-N60 library).

### PAL-seq

Sequencing of endogenous mRNA poly(A)-tail lengths was performed with PAL-seq v3 (for frog oocytes) or PAL-seq v4 (for frog embryos, fish embryos, and mouse oocytes) as described.^41^ When preparing the sequencing libraries of mRNAs from fish embryos, a different 3ʹ adapter (KXS013) was used for 3ʹ-end ligation, and poly(A)-selected mRNA from HeLa cells was used as spike-in, replacing poly(A)-selected mRNA from zebrafish ZF4 cell line normally used as the spike-in. After the first round of sequencing revealed that many of the reads from the fish mRNA libraries were for 5.8S rRNA, the cDNA of 5.8S rRNA was depleted from the cDNA libraries with an antisense oligo as follows. The seven cDNA libraries made from mRNAs of zebrafish embryos at different stages were mixed at roughly equal molar ratios in a total of 5 fmol, and then 50 pmol KXSH015 and 2x SSC were added in a total of 100 µl. The cDNAs and the oligo were annealed by incubation at 65°C for 5 min and then slowly cooled to 23°C at 0.1°C/sec. The annealed mixture was combined with 100 µl MyOne Streptavidin C1 beads and incubated for 20 min at 23°C on a thermal mixer, shaking with 15 sec on and 1 min 45 sec off. The supernatant was separated from the beads with a magnetic rack. The beads were washed once with 200 µl 1xB&W buffer. The supernatant and the wash were combined, precipitated with ethanol, and resuspended in 30 µl water. After this 5.8S depletion, libraries were sequenced again as technical replicates. Sequencing data of replicates were merged for each sample after confirming consistency.

### Genome references and gene annotations

Human (release 25, GRCh38.p7, primary assembly) and mouse (release 10, GRCh38.p4, primary assembly) genomic sequences were downloaded from the GENCODE website (https://www.gencodegenes.org). Sequences of mitochondrial pseudogenes in human and mouse genomes were masked as described.^41^ Frog (*X. laevis*) genomic sequences (v10.1 assembly) were downloaded from the Xenbase website (www.xenbase.org). The frog mitochondrial genomic sequence was obtained from NCBI website (NC_001573.1) and appended to the frog genome. Fish (*Danio rerio*) genomic sequences (GRCz11) were downloaded from the Ensembl website (www.ensembl.org).

Human (release 25, GRCh38.p7, primary assembly) and mouse (release 10, GRCh38.p4, primary assembly) gene annotations were downloaded from the GENCODE website. Frog (*X. laevis*) gene annotations (v10.1 assembly) were downloaded from the Xenbase website. Frog mitochondrial gene annotations were curated based on the information obtained from the NCBI website (NC_001573.1) and appended. Fish (*Danio rerio*) gene annotations (GRCz11.v97) were downloaded the Ensembl website.

For each species, the gene annotation file was processed for downstream analysis with a custom script in the following steps: 1) non-protein-coding annotations were removed; 2) annotation entries without “gene_id” or “transcript_id” were removed; 3) annotation entries (either exon or CDS) of a transcript that overlapped were merged; 4) for each gene, transcript isoforms without CDS annotations were removed if at least one other isoform had CDS annotations, and transcript isoforms without exon annotations were removed if at least one other isoform had exon annotations. The processed gene annotations were referred to as “multi-isoform gene annotations”. The multi-isoform gene annotations for each species were further processed to generate a file referred to as “single-isoform gene annotations”, in which a “main” transcript isoform was annotated for each gene by the following tie-breaker: 1) the isoform with both exon and CDS annotations; 2) the isoform with the most number of exons (only for frog annotations); 3) the isoform that was the longest (summing all exons); 4) the isoform with the longest CDS.

### Annotation of mRNA 3ʹ-end isoforms

PAL-seq tags were used to annotate mRNA 3ʹ-ends because they provided information on cleavage and polyadenylation sites. Software HOMER^81^ was used to call peaks in the “tss” style, with PAL-seq data as the “Test” group and RNA-seq data as the “Control” group, and the following parameters “-style tss - strand separate -fdr 0.001 -ntagThreshold 10 -size 41 -center”. Uniquely mapped reads in same cell type of the same species were merged for both PAL-seq data and RNA-seq data. Specifically, the following datasets were used for the input: 1) frog oocytes/embryos: PAL-seq data for prophase I-arrested oocytes (stage VI) at 0, 1, 3, 5, 7, and 9 h post progesterone treatment (Test) and RNA-seq data for prophase I-arrested oocytes (stage VI) oocytes^41^ (Control); 2) fish embryos: PAL-seq for embryos at 0, 1, 2, 3, 4, 5, and 6 h post fertilization (Test) and RNA-seq data for embryos at 0, 1, 2, 3, 4, 5, 6, 7, and 8 h post fertilization^82^ (Control); 3) mouse oocytes: PAL-seq data for GV and MII oocytes (Test) and RNA-seq data for GV, MI, and MII oocytes^12^ (Control). 4) human oocytes: the 3ʹ-end coverage of the PacBio-based long-read sequencing data from PAIso-seq in human GV oocytes^28^ (Test) and RNA-seq data from GV oocytes^60^ (Control).

A custom script was used to filter peaks and associate each of them with an mRNA isoform of a gene in the following steps: 1) the center position of each peak was extracted as the UTR 3ʹ-end; 2) peaks on the mitochondrial chromosome were removed; 3) *bedtools* (v2.26.0)^83^ *intersect* was used with the “single-isoform gene annotations” to find the mRNA isoform that uniquely overlapped with each peak, and peaks that overlapped with more than one mRNA were removed; 3) for peaks that didn’t overlap with any mRNAs in step 2, *bedtools intersect* was used with the “multi-isoform gene annotations” to find the gene that uniquely overlapped each peak, and peaks that overlapped with more than one gene were removed. In cases in which a peak overlapped with more than one mRNA isoform of the same gene, the mRNA isoform with the longest exon that overlapped with the peak was chosen; 4) for peaks that had not overlapped with any mRNAs in previous steps, *bedtools closest* was used with the “single-isoform gene annotations” to find the closest exon of an mRNA isoform upstream of each peak.

After filtering peaks and associating each of them with an mRNA isoform, a “high” or “low” confidence tag was assigned to each peak for further filtering in downstream analysis based on the following criteria: 1) if a peak overlapped with the last exon of its associated gene, it was labeled as “high” confidence; 2) if a peak was downstream the last exon of its associated gene, a distance between the peak and the last exon was calculated. If this distance was smaller than 500 nt, the peak was labeled as “high” confidence. If this distance was larger than 10,000 nt, the peak was labeled as “low” confidence. 3) for the remaining peaks, a density-ratio test was performed to determine the confidence level of each peak. To perform this test, sequencing-depth normalized RNA-seq read densities were calculated for two regions of each peak. The first region was the exon immediately upstream of the peak (partial exon if the peak and the exon overlapped or full exon if they didn’t overlap). The second region was 150 nt immediately upstream of the peak. The ratio of the read densities of these two regions was calculated for all peaks (including those with a labeled confidence tag). A reference distribution of the ratio values for all peaks that overlapped the last exons of genes was obtained. For each remaining peak without a labeled confidence tag, if the ratio value for the peak was more than 1.5 inter-quantile range (IQR, difference between values at 25^th^ percentile and 75^th^ percentile) above the 75^th^ percentile or less than 1.5 IQR below the 25^th^ percentile of the ratio values of the reference distribution, it was labeled as “low” confidence. Otherwise, it was labeled as “high” confidence. Annotations of mRNA 3ʹ-end isoforms used in this study are provided in Table S2.

### 3ʹ-UTR annotations and sequences

For each organism, mRNA 3ʹ-UTR annotations were curated with a custom script from the “muti-isoform gene annotations” and the “mRNA 3ʹ-end isoform annotations” generated in this study. For each mRNA 3ʹ-end in the “mRNA 3ʹ-end isoform annotations”, if the 3ʹ-UTR annotation of the 3ʹ-end-associated mRNA isoform was available in the “multi-isoform gene annotations”, the 3ʹ-UTR annotation was extracted and the genomic coordinate of the 3ʹ-end was updated from the information in the “mRNA 3ʹ-end isoform annotations”. If the 3ʹ-UTR annotation of the 3ʹ-end-associated mRNA isoform was unavailable, the 3ʹ-UTR annotation was derived in one of the following ways: 1) if both the exon and CDS annotations were available, the exons downstream of the last nucleotide of the CDS were extracted as the 3ʹ-UTR and the genomic coordinate of the 3ʹ-end was updated from the information in the “mRNA 3ʹ-end isoform annotations”; 2) if the CDS annotation was available but the exon annotation was unavailable, the genomic region between the last nucleotide of the CDS and the 3 ʹ-end in the “mRNA 3ʹ-end isoform annotations” was used as the 3ʹ-UTR. If the 3ʹ-end of the curated 3ʹ-UTR had a confidence label “low” in the “mRNA 3ʹ-end isoform annotations,” then it was assigned the label “uncertain,” which was used for filtering low-confidence 3ʹ-UTRs in downstream analyses. For mRNA 3ʹ-ends with 3ʹ-UTRs that could not be annotated as described above because either 1) the 3ʹ-end was upstream of all 3ʹ-UTR annotations; 2) the 3ʹ-end was upstream of the last nucleotide of the CDS, or 3) neither CDS nor 3ʹ-UTR annotations was available, a 2000-nt genomic region upstream the 3ʹ-end was used as the putative 3ʹ-UTR and this 3ʹ-UTR was assigned the label “uncertain”. The curated mRNA 3ʹ-UTR annotations are provided in Table S3.

Sequences corresponding to the mRNA 3ʹ-UTR annotations were extracted from genomic sequences using *bedtools getfasta*. Sequence fragments of each mRNA 3ʹ UTR were stitched together with a custom script. Unless indicated, all “uncertain” 3ʹ UTRs were excluded from downstream analyses.

### PAL-seq data analysis

PAL-seq v3 reads were trimmed with *cutadapt* (v3.7)^84^ with the parameters “-m 15 --quality-base=64 -q 20,20 --match-read-wildcards -e 0.05 -a NNNNATCTCGTATGCCGTCTTCTGCTTG -O 7”. The trimmed reads were mapped using *STAR* (v2.7.1a)^85^ to the reference database containing the genomic sequences of the organism from which the mRNAs were obtained, the genomic sequences of humans or fish, depending on which spike-in RNAs were used, and the sequences of the RNA poly(A)-tail-length internal standards^9^ that had also been added to each sample, with the parameters “--runThreadN 16 -- runMode alignReads --outFilterMultimapNmax 1 --outReadsUnmapped Fastx --outFilterType BySJout -- outSAMattributes All --outSAMtype BAM Unsorted SortedByCoordinate”. Uniquely mapped reads were intersected with a database containing mRNA 3ʹ-end annotations of the organism from which the mRNAs were obtained, the single-isoform gene annotations of humans or fish, depending on which spike-in RNAs were used, and 3ʹ-end annotations of the poly(A) standards, using *bedtools intersect* with the parameters “-wa -wb -bed -s” and retaining reads that corresponded to a unique mRNA isoform. For each library, 10% of reads that intersected with the spike-in RNA annotations (but no more than 50,000 and no less than 5,000) were randomly picked as the training set for determining the Hidden Markov Model (HMM).

To determine the poly(A)-tail length, a T-signal value *S_i,j_* was calculated for each sequencing cycle *i* of read 2 of each read cluster *j* by the following formula:

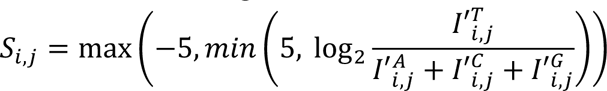

where 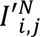 is the normalized intensity of channel N (A, C, G, or T) at cycle *i* of read 2 of cluster *j* defined by the following formula:

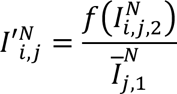

where 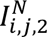 is the raw intensity of channel N at cycle *i* of read 2 of cluster *j* and the function *f* is defined by the following:

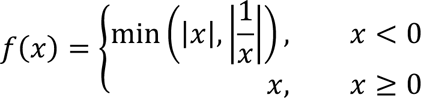

and 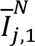 is the average intensity of channel N of read 1, defined by

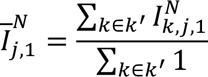

where 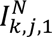 is the raw intensity of channel N at cycle *k* of read 1 of cluster *j*, and *k*’ indicates a set of cycles in the range of 15 to 40 where a base called by the sequencer is the same as the identity of the channel N in read 1 of cluster *j*. For cases in which *S_i,j_* could not be calculated because the normalization of a channel could not be performed or 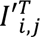 was 0, *S_i’, j_* from neighboring cycles (*i* − 5 ≤ *i*’ ≤ *i* + 5) were averaged to infer *S_i,j_*. If a cluster had more than 5 cycles for which *S_i,j_* values were not available, it was discarded.

A five-state mixed Gaussian Hidden Markov model (from the Python *ghmm* package) was trained on the T-signal values of the training set, and the trained model was used to decode the sequence of the states from the T-signal values of the entire dataset. The initialization parameters and the process of determining poly(A)-tail lengths from decoded states were as described in previous PAL-seq data analyses.^41^

PAL-seq v4 data analysis was performed similarly to PAL-seq v3, with the following differences. First, the *k*’ set used to obtain normalized intensity 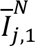 included cycles in the range of 20–50, due to the longer length of read 1 in PAL-seq v4. Second, the poly(A)-tail length was determined by the length of the longest consecutive states that started before cycle 3 and were in the state 1 or 2. Third, for the fish embryo datasets, a different transition matrix was initialized for the HMM to allow backward state transitions:

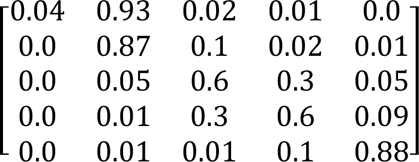

### Reporter 3ʹ-UTR variant and tail-length analysis

For the N60-PAS*^mos^* library, sequencing reads from the FASTQ file were first examined for the expected constant sequence AATAAAGAAATTGATTTGTCT at position 61–81, allowing positional offsets between –3 and +3 nt. Reads with a 21-nt sequence that had no more than six mismatches to the constant sequence at any allowed positions were retained. For each retained read, the sequence preceding the constant sequence was recorded as the 3ʹ-UTR, and the region after this segment was used to determine the poly(A)-tail length. The retained reads were examined for the presence of the 3ʹ-adapter sequence TCGTATGCCGTCTTCTGCTTG with 4 allowed mismatches. If the adapter sequence was identified for a read and the position of the adapter was before the starting position of the poly(A) tail, this read was discarded. If the 3ʹ-UTR of a retained contained an “N” nucleotide or if the lowest quality score of the 3ʹ-UTR was not larger than 10, this read was discarded. One percent of the retained reads (no more than 50,000) were randomly picked as the training set for determining the Hidden Markov Model (HMM).

The poly(A)-tail length was determined similarly to that in the PAL-seq v4, with the following differences: 1) an A-signal value *S*_!,#_was calculated for each sequencing cycle *i* of each read cluster *j* by swapping the A and T channels in the formula for the T-signal value; 2) the 3ʹ-UTR replaced read 1 in the PAL-seq analysis and the entire region was used to determine the average intensity of each channel; 3) the poly(A)-tail length region defined by the constant-sequence segment replaced read 2 in the PAL-seq analysis (cycle 1 was the first cycle after the constant-sequence segment); 4) if a cluster had more than 50 cycles in which *S*_!,#_ values were not available, it was discarded; 5) the transition matrix was initialized as it was used for the fish embryo datasets, allowing backward transitions between HMM states; 6) the poly(A)-tail length was determined by the length of the longest consecutive states that started before cycle 5 and were in the state 1 or 2. Due to the high sequence complexity of each library, most 3ʹ-UTR sequences of these complex libraries were measured only once; for those that were measured more than once, the median value of poly(A)-tail lengths was used.

Analysis of the CPE*^mos^*-N60 library was performed similarly to that of the N60-PAS*^mos^* library, except no read-filtering was performed based on the constant region, because in contrast to the N60-PAS*^mos^* library, this library did not contain a constant region in the sequencing read. Instead, the entire region of the first 60 nt was used as the 3ʹ-UTR, and for determining the poly(A)-tail length, the start of the poly(A) region was set at cycle 61. Sequences of some 3ʹ-UTR variants of this library had strong similarities to the last 50 nt of the coding region of the mRNA reporter or to the 5.8S rRNA. These were presumably sequencing artifacts or contaminants. To exclude them, all 3ʹ-UTR sequences of this library were aligned to the last 50 nt of the Nanoluc sequence and the 5.8S rRNA using the Python package *Biopython*^86^, and those that had an alignment score higher than 25 with the Nanoluc sequence or an alignment score higher than 30 with the 5.8S RNA were removed.

Analysis of the N60 and N37-PAS-N17 libraries was performed similarly to that of the CPE*^mos^*-N60 library. Because these two libraries were mixed for co-injection and had the same 3ʹ-UTR length, their sequencing results were processed together until the last step, at which the N60 and N37-PAS-N17 libraries were separated based on whether the sequence at positions 38–43 matched a PAS (AATAAA).

### *K*-mer-associated tail length and tail-length change of reporter mRNAs

To obtain statistics (mean, median, and standard deviation) of poly(A)-tail length associated with a *k*-mer in an mRNA reporter library dataset, all 3ʹ-UTR variants that contained the *k*-mer within their variable region were counted, and each unique 3ʹ-UTR variant contributed equally to tail-length statistics. When examining the positional effect of a *k*-mer, only variants with a *k*-mer at a specific position were considered. The position of a *k*-mer was determined by its position relative to the 3ʹ-end of the variable region for the N60-PAS*^mos^* library, or to the 5ʹ-end of the variable region for the other three libraries. When examining the effect of the co-presence of two or more *k*-mers, only variants containing the specified *k*-mers at non-overlapping positions were considered. When examining the effect of flanking nucleotides of a *k*-mer, only variants containing the *k*-mer and an indicated flanking nucleotide at the indicated position relative to the *k*-mer regardless of the position of the kmer, were considered.

To examine the tail-length change associated with a *k*-mer between two datasets, the difference was calculated between the two mean values of the tail length associated with the *k*-mer in each dataset. When indicated, a one-sided Welch’s *t*-test was performed with the calculated means, standard deviations, and sample sizes associated with the *k*-mer against a reference value, either 0 or the difference of mean tail lengths of all 3ʹ-UTR variants in the two datasets compared. A Z score of tail-length change was calculated for each *k*-mer with respect to all *k*-mers of the same length.

In some cases, *k-*mer-associated tail-length differences (*k-*mer of the same length) were examined in an iterative exclusion process. In each round of the iteration, mRNA variants with the variable region containing any *k-*mer in the *k*-mer exclusion list (starting with an empty list in the first round) were excluded from the analysis. The average tail-length difference was then calculated for all *k*-mers that were not in the exclusion list. The result of the *k*-mer with the largest tail-length increase (or decrease) was recorded, and this motif was added to the exclusion list before starting the next round. The iteration ended when either there was no *k*-mer motif associated with statistically significant increased (or decreased) tail length (Bonferroni correction-adjusted *P* < 0.05 in a Welsh’s *t*-test) or no *k*-mer motif had a Z score of the tail-length difference higher than 2 (or lower than –2 when tail-length decrease was examined).

### Sequence motifs as binary predictors of endogenous cytoplasmic polyadenylation targets

The tail-length change was compared between the median tail length at 0 h post progesterone and that at 7 h post progesterone for endogenous mRNAs in frog oocytes. Only mRNA isoforms that met the following criteria were included in the analysis: 1) the length of the 3ʹ-UTR was longer than 10 nt; 2) the 3ʹ-UTR not labeled as “uncertain” in the annotations; 3) the 3ʹ-UTR contained a PAS (AAUAAA or AUUAAA) within the last 150 nt of the 3ʹ-end, and 4) the mRNA isoform had more than 50 poly(A) tags in both datasets that were compared. For a given threshold of tail-length difference (ranging from 0–10 nt), mRNA isoforms with tail-length changes larger than the threshold were treated as true positives. The classification threshold was the negative value of the distance *d* to the 3ʹ-end. When a *k*-mer or a *k*-mer group was examined as the binary predictor for tail-length changes, for a given classification threshold −*d* (*d* > 0), an mRNA isoform was considered positive if this *k*-mer or one of the *k*-mers in the *k*-mer group was found within the last *d* nt of the 3ʹ-UTR of this mRNA isoform. Values of true positive rate (TPR) and false positive rate (FPR) were calculated at all possible classification thresholds using the Python *scikit-learn* package^87^ and were used to make the receiver operating characteristic curve (ROC). The area-under-curve (AUC) values were also calculated with the Python *scikit-learn* package^87^.

### Motif logos associated with tail-length changes

Motif logos were made with 8-mers associated with tail-length changes. *k*-mers were filtered with indicated cutoff values for the tail-length difference, the Z score of the tail-length difference, and the Bonferroni correction-adjusted *P* value (Welsh’s *t*-test for tail-length changes). The weights used for producing motif logos were either tail-length changes or Z scores, as indicated. Retained *k*-mers were ranked by the weight in a descending (when motifs associated with tail-length increase were examined) or ascending order (when motifs associated with tail-length decrease were examined). In an iterative procedure starting with the top-ranked *k*-mer, each *k*-mer was aligned to all *k*-mers in a seed list (starting with an empty list), except for the first *k*-mer, which was added to the seed list without any alignments. All alignments were performed without allowing gaps. Alignment scores were calculated for all possible alignment positions between the *k*-mer and all seeds, adopting the following scoring scheme: 1 for a match, –1.5 for a mismatch, –1 for a position offset. The alignment with the highest score (must be higher than –0.1) was kept and the weight was assigned to this *k*-mer-seed alignment. If the highest score was tied among multiple alignments, either between the *k*-mer and the same seed at different positions or between the *k*-mer and different seeds, all tied alignments were kept and the weight was evenly distributed among these alignments. If none of the alignments had a score higher than –0.1, this *k*-mer was considered not aligned and it was added to the seed list. This iterative procedure stopped when all retained *k*-mers had been added to the seed list or had been aligned.

A position weight matrix (PWM) was calculated for each *k*-mer seed in the seed list based on the weights of the seed and all the *k*-mers aligned to it. For each position of the alignments, the weight of each nucleotide contributed by each *k*-mer was summed. If a *k*-mer had no nucleotide aligned at this position, the weight was evenly distributed between A, C, G, and U nucleotides. The summed weights of nucleotides at each position were normalized to the total weights of that position. Motif logos were generated from the resulting PWMs with the R package ggseqlogo^88^.

For the N60-PAS*^mos^* library in frog oocytes, *k-mer*-associated tail-length difference was calculated in an iterative exclusion process. We found this process was computationally heavy and the motif logos generated from this result were similar to those generated from *k*-mer-associated tail-length difference not calculated in an iterative exclusion process. As a result, for all other datasets, *k*-mer-associated tail-length differences used to generate motif logos were not calculated in an iterative exclusion process.

In Figure 1E, tail-length changes were examined in an iterative process. A cutoff of Z score ≥ 3 and tail-length difference ≥ 2 was applied. Tail-length changes were used as weights. In Figure S3G&H, A cutoff of Z-score ≥ 3 (for tail lengthening) or Z-score ≤ –3 (for tail shortening) was applied. Z-scores were used as weights.

### Structural accessibility and tail lengths

The structural accessibility was predicted for each 3ʹ-UTR variant with the entire 3ʹ-UTR sequence (including the variable region and the constant region) using RNAfold^78^ with the parameters “--no PS -- filename-full -p1”. Both the minimum of free energy (MFE) of the 3ʹ-UTR and the base-pairing probability of each base were obtained from the output. The unpairing probability of a motif was the geometric mean of the unpairing probability of each nucleotide of this motif, defined by

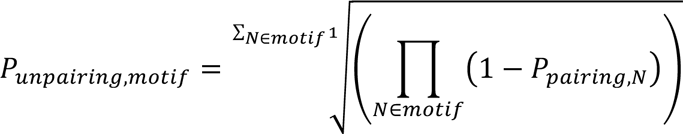

where *P_pairin,N_* is the base-pairing probability of the nucleotide *N* in the motif.

When examining the effect of structural accessibility on tail-length changes for the N60-PAS*^mos^* library during frog oocyte maturation, only variants containing a single CPE (UUUUA) were considered. Tail lengths of variants at 7 h post progesterone treatment were used as a proxy for assessing the tail-length changes from 0 h to 7 h, because most variants were not observed in both time points due to the sequence complexity of the library. In addition, because most variants had initial tail lengths of 35 nt at 0 h post progesterone treatment, the end-point tail lengths reflected tail-length changes. To minimize the potential confounding effect of GC-content of the variant sequence, variants containing a CPE (UUUUA) at indicated positions were divided into 20 equal-sized groups, based on their GC-contents, and the correlation coefficients between the tail length and a structural accessibility metric of interest (MFE or unpairing probability of a motif) were calculated for all the variants within each group.

### Motif-associated tail-length change for endogenous mRNAs

Tail-length changes were calculated for all mRNA isoforms with unique 3ʹ ends by comparing their median tail lengths measured at two different developmental stages, requiring no fewer than the indicated number of poly(A) tags at both stages. To examine the tail-length change associated with a motif, mRNAs were divided into two groups based on whether they contained the motif of interest within their 3ʹ UTRs. When specified, the two groups were subsetted to match the initial tail-length distribution and the 3ʹ-UTR length distribution using the R software package MatchIt^89^ with parameters “distance = ‘glm’, method = ‘cem’, k2k = TRUE”. Tail-length changes of mRNAs from the two groups or the two subsetted groups were used to make cumulative distribution plots.

### Enrichment analysis of *k*-mer-associated translational efficiency

*k*-mer-associcated translational efficiency *E^k^* is defined as:

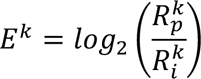

where 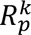 is the fraction of mRNA variants in the library that contained the *k*-mer (or multiple *k*-mers when their co-occurrence was evaluated) in the pellet (for sucrose cushion) or eluate (in the case of nascent-chain pulldown), and 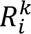 is the fraction of mRNA variants in the library that contained the *k*-mer in the input. 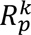 and 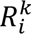 are defined as

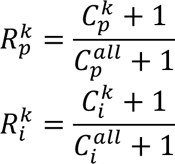

where 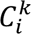, 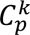 represent the numbers of mRNA variants that contained the *k*-mer in the input and the pellet respectively. 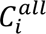 and 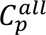 represent the number of total mRNA variants in the input and the pellet respectively. When indicated, a binomial test was performed between 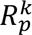 and 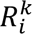 by assuming 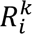 as the expected success probability, 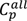 + 1 as the total number of trials, and 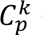 + 1 as the number of successes.

### Measurement of poly(A)-tail lengths in mouse and human oocytes

Poly(A) tail-length measurements by PAL-seq in mouse GV oocytes and MII oocytes were merged for each pair of replicates, after confirming reproducibility. The tail-length change for each mRNA 3ʹ UTR isoform was calculated between the median tail length in GV oocytes and that in MII oocytes, requiring ≥ 50 poly(A) tags in each of the two merged datasets.

For human oocytes, PAIso-seq^28^ bam files were downloaded from Genome Sequence Archive for Human (GSA-Human: https://ngdc.cncb.ac.cn/gsa-human/) with the accession number HRA001911. The reverse-complement sequence of the soft-clipped region at the 5ʹ end of each read was extracted from the bam file. The poly(A)-tail length was determined by the length of the longest consecutive As (allowing individual non-A nucleotides within the stretch) that started before the third nucleotide of the extracted sequence. The tail-length change for each mRNA 3ʹ UTR isoform was calculated as the median tail length in GV oocytes compared to that in MII oocytes, requiring ≥ 50 poly(A) tags in each dataset.

### Tail-length changes of homologous genes in frogs, mice, and humans

The frog-to-human gene ID conversion table was downloaded from Xenbase. The mouse-to-human gene ID conversion table was downloaded from the Ensembl website. Measured tail-length changes were compiled for human MII and GV oocytes, mouse MII and GV oocytes, and frog oocytes 0 and 7 h post progesterone treatment. If a gene had multiple mRNA isoforms, the dominant isoform (> 50% by tag count in GV oocytes of humans and mice or frog oocytes 0 h post progesterone treatment) was chosen to represent that gene. If more than one homolog of a human gene was identified in another species (mice or frogs), the mean tail-length change of the homologs was used.

### Relationship between poly(A)-tail length and TE in human and mouse oocytes

For mouse oocytes, TEs measured by ribosome footprint profiling and mRNA-seq were obtained from a published study^12^, and genes were filtered to require TE > 0 in both GV and MII oocytes. For human oocytes, normalized read counts of ribosome footprint profiling and mRNA-seq were obtained from a published study^60^, and TEs were calculated for genes that had a normalized read count of ribosome protected fragments > 0 and normalized read count of mRNA-seq tags > 1 in both GV and MI oocytes. For both mouse and human oocytes, mRNA isoforms were required to have ≥ 50 poly(A) tags in each dataset for calculating tail-length changes. If a gene had more than one mRNA isoform, only the one with ≥ 90% of the total poly(A) tags of that gene was retained.

### Data and code availability

Raw and processed sequencing data were deposited in the GEO database (GSE241107). PAL-seq data analyses were performed using a custom script written in Python 2.7 and available at https://github.com/coffeebond/PAL-seq. Reporter mRNA tail-length sequencing data analyses were performed using a custom script written in Python 3.6 and available at https://github.com/coffeebond/MPRA_tail_seq.

